# Meiotic Double-Strand Break Proteins Influence Repair Pathway Utilization

**DOI:** 10.1101/376582

**Authors:** N. Macaisne, Z. Kessler, J.L. Yanowitz

**Affiliations:** Magee-Womens Research Institute, Dept. of Obstetrics, Gynecology, and Reproductive Sciences, University of Pittsburgh School of Medicine, Pittsburgh PA 15213

**Keywords:** DNA repair, Meiosis, Double-strand break, *C. elegans*, Pathway choice

## Abstract

Double-strand breaks (DSBs) are among the most deleterious lesions DNA can endure. Yet, DSBs are programmed at the onset of meiosis and are required to facilitate appropriate reduction of ploidy in daughter cells. Repair of these break is tightly controlled to favor homologous recombination (HR), the only repair pathway that can form crossovers.

However, little is known about how the activities of alternative repair pathways are regulated at these stages. We discovered an unexpected synthetic interaction between the DSB machinery and strand-exchange proteins. Depleting the *C. elegans* DSB-promoting factors HIM-5 and DSB-2 suppresses the formation of chromosome fusions that arise in the absence of RAD-51 or other strand-exchange mediators. Our investigations reveal that non-homologous and theta-mediated end joining (c-NHEJ and TMEJ, respectively) and single strand annealing (SSA) function redundantly to repair DSBs when HR is compromised and that HIM-5 influences the utilization of TMEJ and SSA.

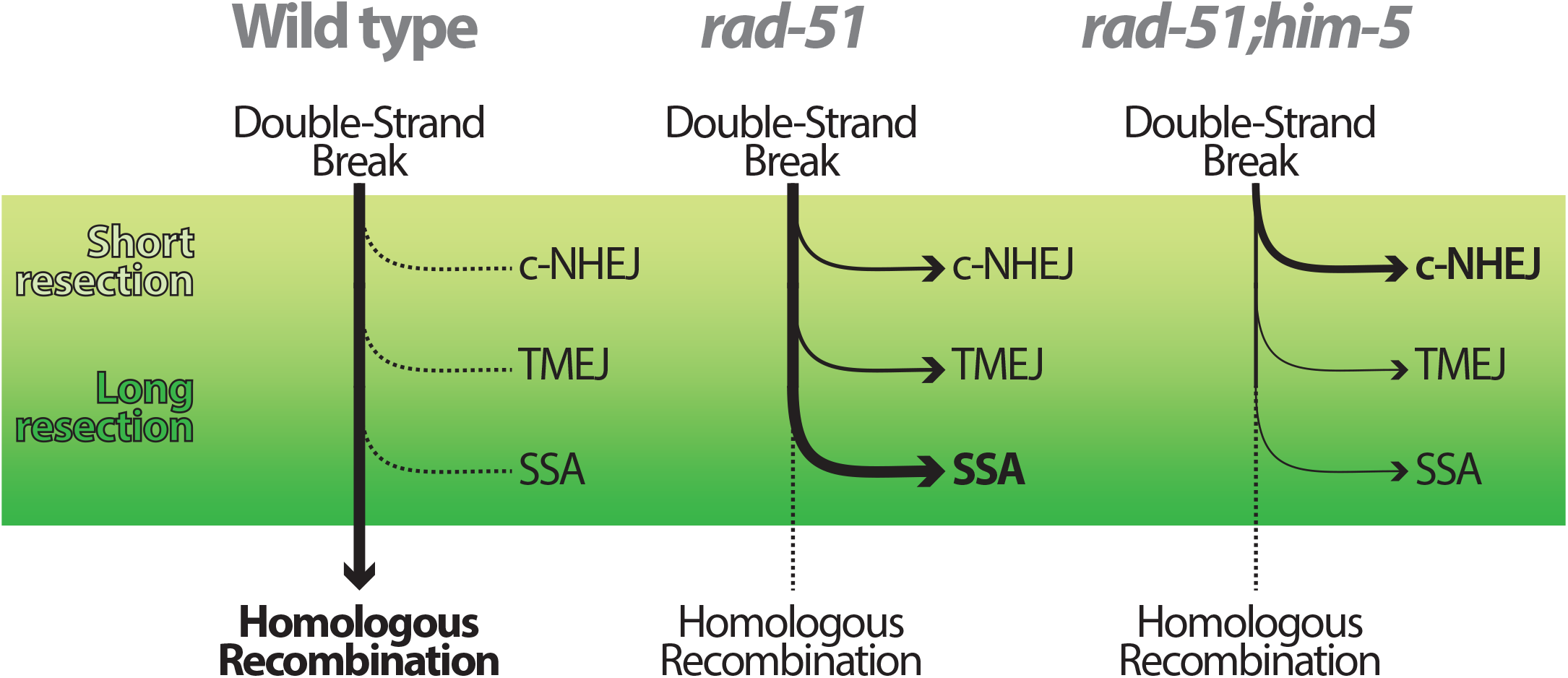

## INTRODUCTION

DNA is under constant stress from endogenous and exogenous sources. The resulting lesions must be repaired to ensure genome integrity. In the germ line, improperly repaired damage can lead to inherited mutations or chromosomal rearrangements, which are often associated with heritable predispositions to cancer, recurrent miscarriage, and genetic disorders. Double-strand breaks (DSBs) are considered the most toxic type of lesion to the genome and several pathways exist to repair them. The mechanism of repair depends on DSB structure and cell cycle phase (Shrivastav *et al*. 2008; Chang *et al*. 2017).

Cells in S and G2 phases primarily repair DSBs by homologous recombination (HR), which relies on the presence of a homologous template (the attached sister chromatid or the homologous chromosome) to ensure the most accurate repair. Alternatively, canonical non-homologous end-joining (c-NHEJ) occurs by direct ligating the blunt ends of the DSB without using DNA sequence homology (Branzei and Foiani 2008). Cells leverage c-NHEJ throughout the whole cell cycle (G1, S, G2). Repair by c-NHEJ may generate mutations or chromosome rearrangements, but it can also be conservative if no base is lost after DSB formation (Eid *et al*. 2010). Although it seems that c-NHEJ machinery may be involved in the repair of more intricate DSB structures (Lobrich and Jeggo 2017), other DSB repair pathways are error-prone by nature because they function by annealing sequence repeats. Alternative end-joining (alt-EJ, also called microhomology-mediated end-joining) relies on microhomologies, generally shorter than 10 nucleotides, to ligate cohesive DSB ends, and results in DNA insertions and deletions (Ma *et al*. 2003; Martin *et al*. 2005; Corneo *et al*. 2007; Yan *et al*. 2007; Sakuma *et al*. 2016). Theta-mediated end-joining (TMEJ) is a subtype of alt-EJ that depends on the polymerase theta (polθ). TMEJ can anneal sequences sharing as few as one nucleotide of homology at the junction (Schendel *et al*. 2016). Similarly, but mechanistically independent, single-strand annealing (SSA) can anneal longer sequence repeats (>20 nt) (Maryon and Carroll 1991; Fishman-Lobell *et al*. 1992). Thus, SSA and TMEJ can cause mutations and may be involved in the appearance of more complex chromosome rearrangements (reviewed in (Bhargava *et al*. 2016)).

Despite their potential toxicity, programmed DSBs occur at the onset of meiotic prophase I to induce the formation of crossovers (COs). COs are reciprocal exchanges of genetic material that must occur between each pair of homologs. They form a physical link, the chiasma, which is essential to prevent chromosome nondisjunction. Meiotic DSBs are catalyzed by the conserved and meiosis-specific topoisomerase-like protein SPO11 (Keeney *et al*. 1997).

They are repaired preferentially by HR because it is the only repair pathway that forms COs to guarantee genomic integrity in the resulting gametes. Accordingly, mechanisms must exist in the germ line to ensure that HR predominates over other repair pathways. While many factors promoting HR have been described, it remains poorly understood how alternative repair pathways are suppressed in the germ line (Ranjha *et al*. 2018).

When HR is impaired, c-NHEJ has a well-demonstrated role in meiotic DSB repair (Clejan *et al*. 2006). More recently, TMEJ was shown to participate in DNA repair in *C. elegans* germ cells, although the timing of this repair is unknown (Schendel *et al*. 2016). Notably, SSA has been invoked in repair of DSBs on the X chromosome of hemizigous male nematodes (Checchi *et al*. 2014). Further investigation is needed to decipher how meiotic cells restrict TMEJ and SSA activities. Additionally, although it is known that c-NHEJ predominates in HR-deficient germ lines, it remains unclear whether c-NHEJ competes with HR, coordinates with HR machinery, or simply operates when HR is compromised (Clejan *et al*. 2006).

Recent studies of HR-impaired animals indicate that c-NHEJ is restricted early during DSB processing. DSB end resection starts by nicking the DNA surrounding the catalyzed lesion to remove SPO-11, which is mediated by the MRN complex (MRE-11, RAD-50, NBS-1) endonucleolytic subunit MRE-11 and COM-1 (the worm homolog of the tumor suppressor CtIP/Ctp1/Sae2). The liberated SPO-11 stays covalently attached to small DNA oligos, leaving ssDNA around the DSB site. Further end resection generates a ssDNA template suitable for binding the RecA-like recombinases, RAD-51 or its meiotic-specific paralog DMC1 (Ma *et al*. 2015). In the absence of *com-1*, the Ku-complex (encoded by *cku-70* and *cku-80* in worms) impedes extended resection mediated by the exonuclease EXO-1, and allows for end-to-end ligation at the DSB site by c-NHEJ (Lemmens *et al*. 2013). It has been proposed that meiotic inhibition of c-NHEJ is a result of a competition between MRE-11/COM-1 and the Ku-complex for the access to the DSB site. SPO-11 clipping also allows the access to EXO-1, which probably further prevents Ku binding (Lee *et al*. 1998; Zhou *et al*. 2014). It thus appears that early DSB-processing events coordinate to promote HR over c-NHEJ.

The long single-stranded overhangs are bound by Rad51 and Dmc1 proteins which drive strand invasion and the search for extended sequence homology. In worms, this strand-exchange activity relies on *rad-51*, the only RecA homolog found in this species (Takanami *et al*. 1998). Subsequent repair using the homolog as a template eventually leads to formation of COs. Interestingly, the process of resection also produces DNA intermediates that are compatible with alternative repair pathways, including TMEJ and SSA. However, the role and inhibition of these repair pathways in meiosis remain poorly understood and mostly unexplored.

We report the DSB-promoting factors *him-5* and *dsb-2* promote HR-mediated repair. Depleting either of these genes suppresses chromosomal fusions that occur in *rad-51* mutant germ cells, without abrogating the formation of DSBs. This suppression leaves chromosomes strikingly intact, suggesting that repair occurs robustly through alternative pathways. Through analysis of the *rad-51* and *rad-51;him-5* double mutants, we provide evidence that in addition to c-NHEJ, TMEJ and SSA act as back-up repair mechanisms within the meiotic germ line of HR mutants. Further, we found that *him-5* appears to restrict utilization of SSA and TMEJ, suggesting that meiotic repair pathway choice may be dictated by the DSB machinery itself in order to couple DSB formation to downstream repair via HR.

## MATERIALS AND METHODS

### Strains and growth conditions

*C. elegans* strains were maintained at 20°C under standard growth conditions (Brenner 1974). The N2 Bristol strain was used as the wild-type background (Sulston and Brenner 1974). The strains and genotyping conditions are provided in Table S1 and Table S2, respectively. All strains will be provided upon request.

### CRISPR/Cas9 Mutagenesis

We produced a complete knock-out of the *him-5* gene using CRISPR/Cas9 mutagenesis. Removal of the endogenous *him-5* locus was performed by injecting two gRNAs directed towards NGG motifs located on each end of the him-5 isoform A coding sequence (NM-HIM5sgRNA5 5’-GTCGTTATTAGAACGAATTC-3’ targeting a cut site in exon 1, and NM-HIM5sgRNA6 5’-ATGCGGAATGACCACCAGGC-3’ targeting a cut site in exon 7) with a repair template that bridged the gap with homology arms (JY-NM-086 5’-CCGAAAAAGCAT ACTTAT CATCTGGTTTTTTTTGCTGAAAAT GTC(A)AGAA_CATT(A)CAA AAAAACGCGCTCAATAAttcgtggtttaataatagtagtttt-3’, the gap is represented as an underscore. C to A mutations, shown between brackets were designed in the repair template to avoid Cas9 activity upon successful mutagenesis). The injection was performed according to (Paix *et al*. 2015). We obtained and have analyzed one allele *him-5* that lacks the complete coding sequence *him-5(ea42*) (Table S1).

### Sample preparation and imaging

L4 larvae were aged for 20 hr (day 1 adults) or 47-50 hours (Day 2 adults) before fixation and staining. For Irradiated samples, day 1 adults were exposed to 10 Gy of ionizing radiation using a ^137^Cs source (Gammacell 1000 Elite; Nordion International). Irradiated animals were further aged for 27 hr (day 2 adults) prior to fixation and staining to assess diakinesis phenotypes.

Worm were fixed in Carnoy’s fixative solution (three parts absolute ethanol; two parts chloroform; one part glacial acetic acid), stained with DAPI (4’,6-diamidino-2-phenylindole) for at least fifteen minutes, mounted in Prolong Gold Antifade Mountant with DAPI (ThermoFisher Scientific, P36931), and cured overnight prior to imaging. All images were acquired with a Nikon A1R inverted confocal microscope system, driven by the NIS-elements Software as 0.2 μm Z-stacks. 3D-reconstructed acquisitions were analyzed and captured for publication with the Volocity software (PerkinElmer). Captured images were cropped and assembled using Adobe Photoshop and Illustrator.

### Analysis of DNA structures in diakinesis oocytes and related statistics

DAPI-body content of the −1 nuclei were analyzed individually on 3D reconstructed images in the Volocity software which allows for the free rotation of nuclei. A DAPI-body was defined as a continuous DAPI-stained chromatin structure, independent of its size. A chromatin fragment was defined as a DAPI-body with an estimated volume less than 25% of a univalent observed in *spo-11* mutant −1 nuclei.

The number of DAPI-stained bodies per nucleus was plotted and analyzed with the Graphpad Prism software. We used non-parametric tests as we could not assume ours samples would follow a normal distribution, especially samples with fusions and fragments. We compared the distribution of different DAPI-stained bodies counts (ranks). Pairwise comparisons were done with two-tailed Mann-Whitney tests, with the null hypothesis (H0) that it is equally likely that a randomly selected value from one sample will be less than or greater than a randomly selected value from the second sample, with an error risk of 5%. Multiple comparisons were processed as well with Kruskal-Wallis tests corrected to control the false discovery rate (FDR) using the two-stage step-up method of Benjamini, Krieger and Yekutieli (Benjamini *et al*. 2006). H0 is that the independent samples tested are indistinguishable from the combined population of samples. P-values and q-values (FDR-adjusted P-value) are given in Table 3. H0 was rejected when P was below 0.05 with a FDR lower than 5%. We graphed the median and the interquartile range which are more representative for how each population is distributed. However, we state the average number of DAPI-bodies per nucleus in the text as we thought it is more meaningful to the reader. Summary of statistical tests performed for each figure is provided in Table S3.

## RESULTS

### *him-5* mutations suppress chromosome fusions formed by loss of *rad-51*

In *C. elegans*, defects in meiotic DNA repair can be seen in DAPI-stained nuclei of the diakinesis oocyte positioned just prior to fertilization (−1 nucleus). The morphology and number of DAPI-stained bodies is a readout of crossover formation, with wild-type worms exhibiting 6 ovoid, DAPI-stained bodies corresponding to the 6 pairs of homologs attached by chiasmata (also referred to as 6 bivalents) (Figure 1A, F; (Villeneuve 1994)). When DSB formation is abrogated, the homologous chromosomes dissociate from one another and appear as smaller, round univalents (Dernburg *et al*. 1998). We observe an average of 10.8 DAPI bodies in *spo-11* mutant nuclei, corresponding to a majority of univalent chromosomes, and an occasional bivalent that forms possibly as a consequence of ectopic breaks (Figure 1B, F; see also (Machovina *et al*. 2016). Combinations of univalents and bivalents are seen in mutants with impaired break capability, such as *him-5* and *dsb-2* (Figure 1C, F; Figure Figure S1) (Meneely *et al*. 2012; Rosu *et al*. 2013). In contrast, in mutants defective in strand-exchange activity, the DAPI-stained bodies appear irregularly shaped, often with aggregates (chromosomal fusions) and/or small fragments. For example, the disruption of *rad-51* causes chromatin to form massive clumps (Takanami *et al*. 1998) with an average of 4.23, irregularly shaped and sized DAPI-stained bodies (Figure 1D, F). This phenotype is reminiscent of chromosome end-to-end fusions and is thought to result from random religation of SPO-11 cut chromosomes. Accordingly, these structures are completely dependent on the formation of meiotic DSBs, as seen by their suppression when *spo-11* is co-depleted (Rinaldo *et al*. 2002; Takanami *et al*. 2003).

**Figure 1.**
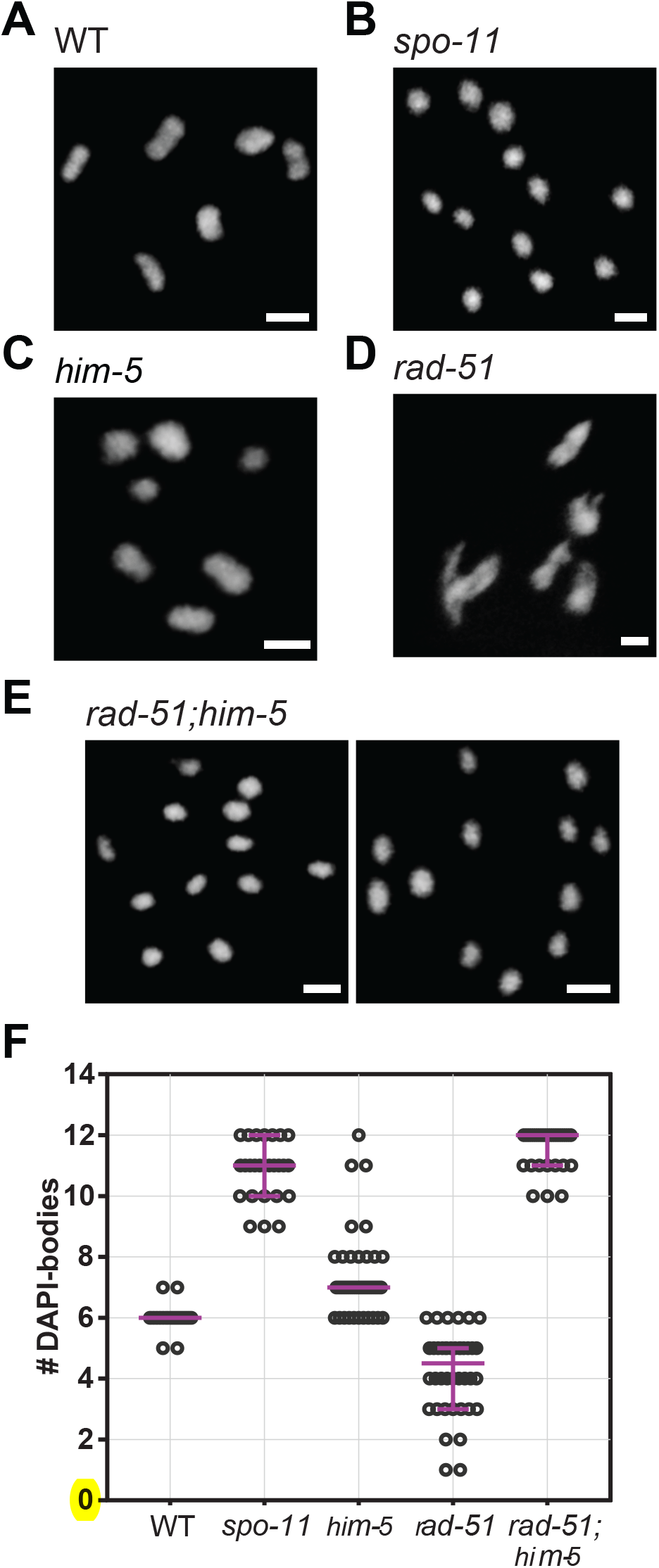
The *rad-51;him-5* double mutant phenocopies *spo-11* with 12 univalent-like structures in diakinesis nuclei. (A-E) Representative images of DAPI-stained diakinesis nuclei of wild type and mutants at day two of adulthood. Scale bars= 2*μ*m. (A) Wild type (n=45); (B) *spo-11(me44*) (n=26); (C) *him-5(e1490*) (n=66); (D) *rad-51(lg8701*) (n=40), (E) *rad-51(lg8701);him-5(e1490*) (n=33). (F) Quantification of DAPI-bodies in diakinesis nuclei. Magenta bars demarcate the median and interquartile range.

We produced the *rad-51;him-5* double mutants for which we expected an intermediate phenotype, with a mixture of chromosome clumps due to the abnormal repair in absence of *rad-51*, and intact X-chromosome univalents that would not have received a DSB due to loss of *him-5* function (Meneely *et al*. 2012). Surprisingly, we instead observed that *rad-51;him-5* diakinesis nuclei contained ~12 DAPI-positive bodies. Moreover, chromatin structures were indistinguishable from intact univalent chromosomes such as those seen in *spo-11* mutants (Figure 1E, F). This is surprising because the *him-5* mutation alone does not prevent DSB formation (Meneely *et al*. 2012). Suppression of *rad-51* chromosomal fusions was observed in *rad-51;him-5(ok1896), rad-51;him-5(e1490*), and *rad-51;him-5(ea42*) (Figure S2), arguing that the suppression is not allele-specific. We thus conclude that loss of HIM-5 function suppresses chromosome fusions formed in *rad-51* mutant animals.

### DSBs are formed and repaired in *rad-51;him-5* mutants

We reasoned that the presence of intact univalents in *rad-51;him-5* could be explained in two ways: either RAD-51 contributes to DSB formation (either directly or via positive feedback); or, HIM-5 prevents the use of alternative DSB repair pathway(s) that can preserve the structure of (univalent) chromosomes. To address whether RAD-51 has a role in DSB induction, we first asked whether the univalents observed in *rad-51;him-5* were specific to loss of *rad-51* or whether depletion of other HR-promoting factors were suppressed by *him-5*. These factors include the RAD-51 cofactor RAD-54 (Ward *et al*. 2010) and the MRX/N complex subunit MRE-11 (Chin and Villeneuve 2001). Although MRE-11 has been shown to promote both DSB formation and DSB end resection, resulting in 12 univalents in null mutant diakinesis nuclei (Chin and Villeneuve 2001), we used separation-of-function allele *mre-11(iow1*) (referred to as *mre-11S* hereafter) which is deficient for end resection, but (mostly) proficient for DSB formation (Yin and Smolikove 2013). In the *mre-11S,him-5* and *rad-54;him-5* double mutants, we observed that, similar to *rad-51;him-5*, diakinesis nuclei contained ~12 DAPI-stained, univalent-like structures, contrasting with the clumps observed in diakinesis nuclei of *rad-54* and *mre-11S* single mutants (Figure 2A, B). Suppression of fusions by *him-5* was also observed in the strand-exchange defective mutants *rfs-1;helq-1* and *brc-2* (Figure S3 A, B). The suppression of multiple early HR factors argues against a role for all of these factors in DSB formation.

**Figure 2.**
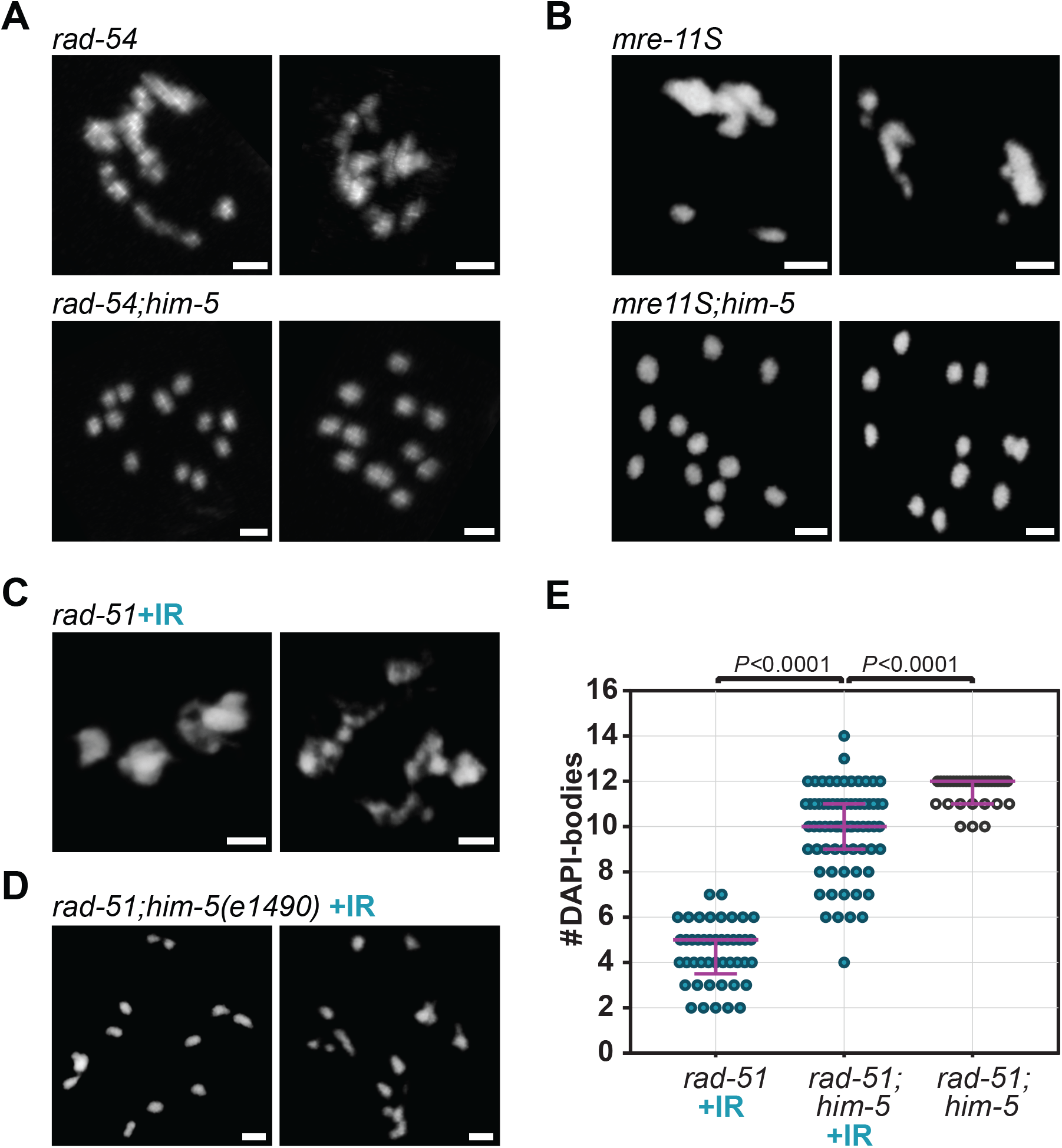
Depletion of *him-5* suppresses chromosome fusions that arise in multiple early HR-defective mutants. (A) *rad-54* and (*B) mre-11S* mutant animals exhibit chromosomal clumping that is suppressed by the depletion of *him-5*. (C, D) Irradiation of *rad-51;him-5* does not restore a *rad-51-like* phenotype when *him-5* is absent (C) *rad-51* + IR control; (D) *rad-51; him-5(e1490*). DAPI-stained diakinesis nuclei at day 1 (A, B) or day 2 (C, D) of adulthood. Scale bars= 2*μ*m. (E) Quantification of DAPI-bodies in diakinesis nuclei of irradiated (+IR; blue dots) *rad-51(lg8701*) (n=45), *rad-51(lg8701);him-5(e1490*) (n=71), and non-irradiated (clear dots) *rad-51(lg8701);him-5(e1490*) (n=33). Magenta bars indicate the median and the interquartile range. P-values calculated with two-tailed Mann-Whitney tests.

To further test whether meiotic DSBs are made in the *rad-51;him-5* mutants, we relied on the fact that chiasma formation can be restored in *spo-11* mutants upon exposure to gamma irradiation (IR) (Dernburg *et al*. 1998). A dose of 10 Gy is expected to induce approximately 20 DSBs and is sufficient to recover at least 1 chiasma per chromosome pair (Yokoo *et al*. 2012; Machovina *et al*. 2016). If *rad-51;him-5* were lacking DSBs entirely, one would expect that 10Gy IR would lead to chromosome fusions, as seen in *rad-51* single mutants. Contrary to this expectation, IR of *rad-51;him-5* failed to induce the *rad-51* phenotype (Figure 2C, D). Instead, irradiated diakinesis nuclei looked remarkably like the unirradiated *rad-51;him-5* with the addition of a few chromosome fusions and occasional chromatin fragments (Figure 2D; Figure S3C). The number of DAPI-stained bodies remained strikingly higher than in irradiated *rad-51* mutants (9.87 vs 4.42 DAPI-bodies/nucleus on average), and closer to the non-irradiated *rad-51;him-5* double mutant (11.61 DAPI-bodies/nucleus). We also evaluated the distribution of DAPI-body counts since in the presence of abnormal chromosome structures, the average can be misleading because outliers can dramatically affect the outcome. In this case, irradiated *rad-51;him-5* nuclei were clearly distinguishable from irradiated *rad-51* nuclei (Figure 2E, Figure S3D; *P*<0.0001), as well as from unirradiated *rad-51;him-5* nuclei (Figure 2E; *P*<0.0001, two-tailed Mann-Whitney tests). Similar results were observed with *mre-11S,him-5:* irradiation did not recapitulate the fusions observed in *mre-11S* single mutants (Figure S3E-G). The failure of IR-induced damage to lead to *rad-51-like* chromosome fusions is consistent with the hypothesis that loss of *him-5* function alters repair outcomes even in the presence of exogenous DSBs.

While the failure of IR to restore fusions in *rad-51;him-5* suggests DSBs can be repaired by alternative mechanisms, this result did not preclude the other possibility that SPO-11-induced breaks were not formed. We thus wanted an alternative approach to verify that SPO-11-mediated DSBs are made in *rad-51;him-5* mutant germ cells. Since there are no direct methods to monitor DSBs (worms do not have H2AX; nor do worms have meiotic crossover hotspots that can be physically monitored for DNA cleavage), we instead took advantage of a chromosome fragmentation assay that requires meiotic DSBs. REC-8 is a meiosis-specific component of a cohesin complex that promotes sister chromatids cohesion. In *rec-8* mutants, DSBs are made, but sister chromatids separate from one another prior to repair, leading to chromosome fragmentation (Pasierbek *et al*. 2001). This fragmentation is dependent on meiotic DSBs as it can be suppressed by co-depletion of *spo-11* (Figure 3A) (Pasierbek *et al*. 2001). Diakinesis nuclei of *spo-11,rec-8* double mutants contained an average of 21.27 DAPI-bodies, close to the expected 24 DAPI-bodies that correspond to the 12 pairs of unattached sister chromatids (The difference between the observed and expected DAPI-body counts might be explained by the inability to unambiguously distinguish closely apposed chromatids). These DAPI-bodies were well formed and uniform in appearance and no chromosome masses or chromatin fragments were observed (n=44; Figure 3A, C), consistent with the lack of DSBs in the absence of *spo-11* function. By contrast to *spo-11;rec-8*, diakinesis nuclei of *rad-51,rec-8;him-5* triple mutants contained an average of 20.54 DAPI-bodies of irregular sizes and shapes (Figure 3B). Greater than 60% of cells contained at least one DNA fragment (n=35; Figure 3B, C), revealing that DSBs are formed in *rad-51;him-5* mutant germ cells.

**Figure 3.**
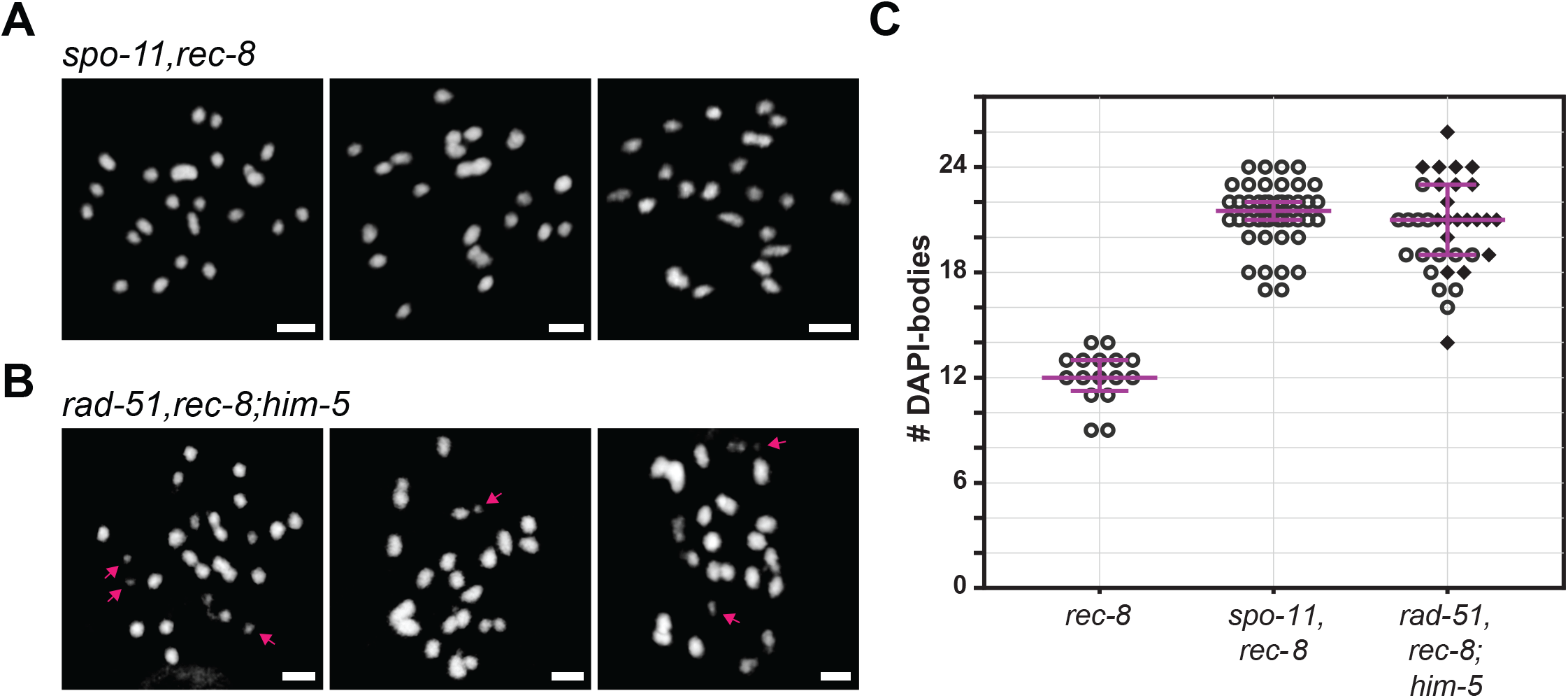
DSBs are observed in *rad-51;him-5*. (A) DNA fragments are not observed in the control *spo-11,rec-8* strain. (B) The presence of DSBs catalyzed by SPO-11 in *rad-51;him-5* is revealed by the presence of chromatin fragments in *rad-51,rec-8;him-5*. Representative images of DAPI-stained diakinesis nuclei in one-day-old adults. Arrows point to chromatin fragments. Scale bars= 2*μ*m. (C) Quantification of DAPI-bodies in diakinesis nuclei: *rec-8(ok978*) (n=16), *spo-11(me44),rec-8(ok978*) (A; n=44), and *rad-51(lg8701),rec-8(ok978);him-5(e1490*) (B; n=35). Diamonds indicate nuclei containing at least one fragment of chromatin. Magenta bars show the median and the interquartile range.

Overall, the presence of meiotic DSBs in *rad-51;him-5* argues that alternative repair pathway(s) must be active in the *rad-51;him-5* germ lines to allow for preservation of chromosome morphology without the formation of chiasmata or fragmentation. We thus hypothesized that *him-5* could influence one or several alternative repair pathways such as c-NHEJ, TMEJ and SSA. However, the involvement of all of these pathways during meiosis, even as backup mechanisms when HR is not available, remains obscure or unexplored.

### Multiple backup DSB repair pathways are active in the germline

It is well documented that c-NHEJ functions in meiosis when strand-exchange activity is depleted (Martin *et al*. 2005; Smolikov *et al*. 2007a; Yin and Smolikove 2013; Mateo *et al*. 2016). In mutant backgrounds where HR is impaired by the loss of *rad-51* or strand exchange activity, chromatin clumps are observed in diakinesis nuclei. These clumps/fusions rely, at least partially, on c-NHEJ as evidenced by the increase in number of DAPI bodies in diakinesis nuclei of *cku-70;rad-51* vs. *rad-51* (Martin *et al*. 2005). We confirmed the role of c-NHEJ activity in meiotic repair in wild type and HR-impaired animals by again quantifying DAPI bodies in −1 diakinesis oocytes. Consistent with prior studies (Clejan *et al*. 2006), we observed that depletion of c-NHEJ did not have a visible effect on meiotic crossover formation: diakinesis nuclei in *cku-80* or *cku-70* mutant animals contained predominantly 6 bivalent-shaped DAPI-bodies, which were indistinguishable from wild-type worms (Figure S4A, B; two-tailed Mann-Whitney tests, P>0.34). When HR function was impaired, chromosome fusions ensued with an average of 4.49 DAPI-stained bodies in *rad-51*. Removal of c-NHEJ functions in this background led to a partial suppression of these fusions with *cku-70;rad-51* and *cku-80;rad-51* exhibiting an average of 5.38 and 5.31 DAPI bodies, respectively, a significant increase in DAPI-bodies compared to *rad-51* (Figure 4A, B; Figure S4C; Kruskal-Wallis, *P*<0.0050). Despite the increased number of chromatin masses, the overall chromatin morphology of these bodies was not qualitatively different (Figure 4A, B). Of note, chromatin fusions were prevalent and chromosome fragments were rare in *cku-70;rad-51* and *cku-80;rad-51*, leading us to posit that multiple pathways contribute to repair.

**Figure 4.**
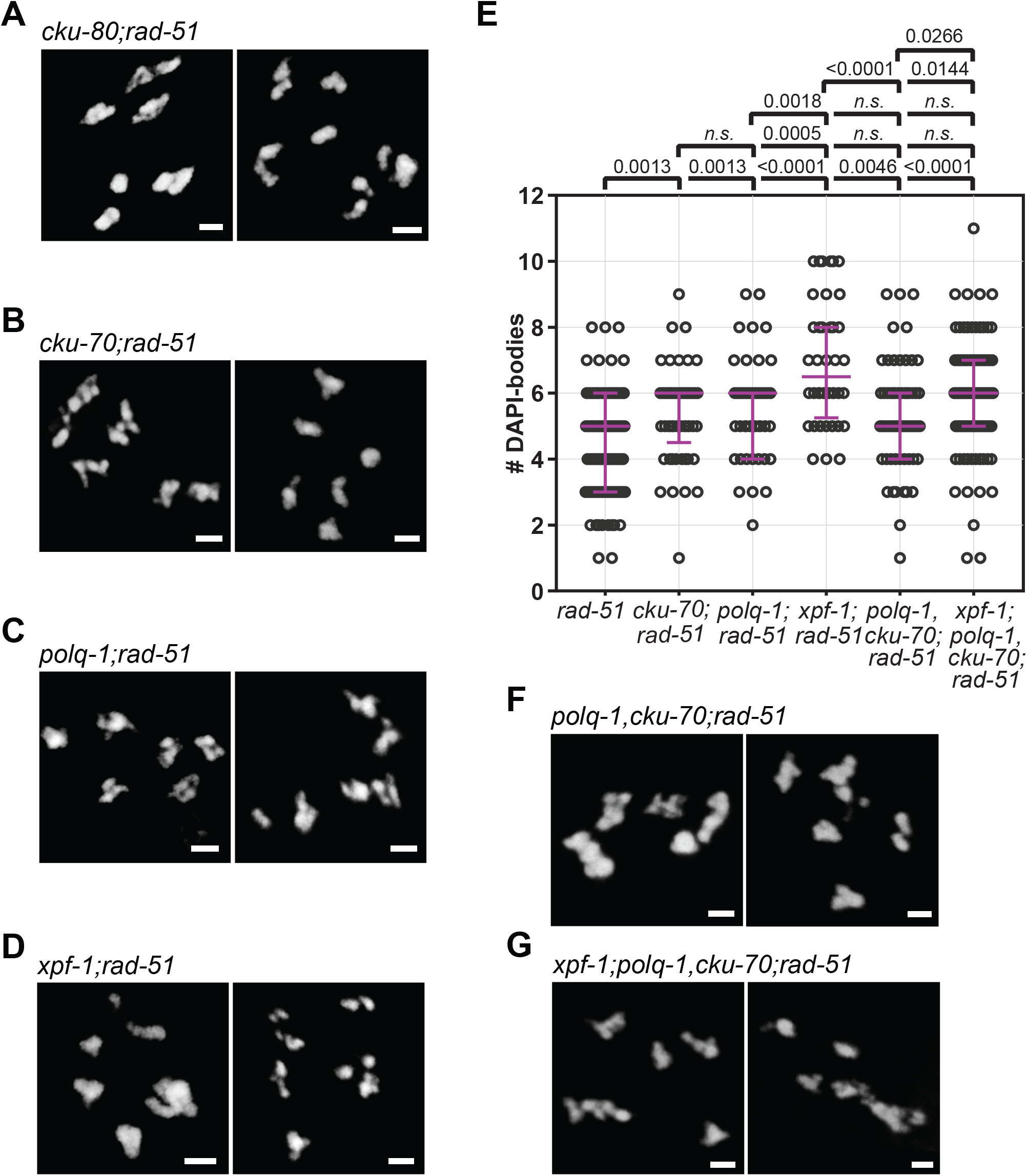
c-NHEJ, TMEJ and SSA contribute to DSB repair in the HR defective *rad-51* mutant (A-D, F, G) Representative images and (E) Quantification of DAPI-stained diakinesis nuclei from one-day-old adults of designated genotypes. Scale bars= 2*μ*m. Magenta bars indicate the median and the interquartile range. P-values are calculated with Kruskal-Wallis multicomparison tests (n.s.: not significant, P>0.19). *rad-51(lg8701*) (n=106), *cku-70(tm1524);rad-51(lg8701*) (n=53), *polq-1(tm2026);rad-51(lg8701*) (n=43), *xpf-1(e1487);rad-51(lg8701*) (n=40), *polq-1(tm2026),cku-70(tm1524);rad-51(lg8701*) (n=60), *xpf-1(e1487);polq-1(tm2026),cku-70(tm1524);rad-51(lg8701*) (n=108).

Although the specifics of meiotic DSB formation and processing have not been fully characterized in worms, short- and long-range resection by MRN/COM-1 and EXO-1, respectively, can generate strand-intermediates that are compatible with error-prone repair pathways known to be active in somatic cells (Truong *et al*. 2013). TMEJ-mediated repair necessitates the annealing of short homology tracks (1-15 bases) to repair DNA, and SSA needs longer tracks of homology (40-300 bases) to function (Sugawara *et al*. 2000; McVey and Lee 2008) reviewed in (Lemmens and Tijsterman 2011)). We set out to determine if TMEJ and SSA are contributing to repair when HR is impaired. To test the involvement of TMEJ, we analyzed mutants in *polq-1*, the worm homolog of the gene encoding polymerase theta (Schendel *et al*. 2016). During meiosis, depleting *polq-1* function did not overtly impair CO formation as revealed by the presence of 6 bivalents in diakinesis nuclei (Figure S4 D, F; Mann-Whitney test, P>0.86). In contrast, *polq-1* mutations significantly increased the average number of DAPI-bodies per cell in *the rad-51* mutant background (5.51 in *polq-1;rad-51* vs 4.49 in *rad-51;* Kruskal-Wallis, P=0.0013) (Figure 4C, E). This tendency to form fewer end-to-end fusions suggests that TMEJ, like c-NHEJ, contributes to repair, seen as chromosomal clumps when HR is defective.

To test the involvement of SSA in the absence of *rad-51*, we analyzed the effects of mutations in *xpf-1*, which encodes the worm homolog of the XPF/Rad1 nuclease. In *C. elegans*, XPF-1 has two described functions: first, as a key contributor to SSA in mitotic cells (Pontier and Tijsterman 2009); second as a component of a meiotic CO resolvase complex (Saito *et al*. 2009). The latter function leads to a defect in chiasmata formation, which we indeed observed in *xpf-1* single mutants (Figure S4 E, F). Since RAD-51 filament formation precedes CO resolution, we hypothesized that any observed effect of *xpf-1* depletion in *rad-51* mutants should be attributed to its role in SSA. As in *cku-70;rad-51* and *polq-1;rad-51*, chromosome morphology was abnormal in *xpf-1;rad-51* (Figure 4D) and significantly less clumping occurred compared to *rad-51* mutants (6.93 vs. 4.49 DAPI-bodies per nucleus on average, in *xpf-1;rad-51* and *rad-51*, respectively, Kruskal-Wallis test, *P*<0.0001) (Figure 4E). The marked increase in DAPI-body numbers (*i.e*. decrease in fusions), even compared to *cku-70;rad-51* and *polq-1;rad-51* (Figure 4E; Kruskal-Wallis, *P*<0.002), suggests that SSA may play a more prominent role than c-NHEJ or TMEJ in DSB repair when HR is defective. This was further confirmed by the co-depletion of alternative repair pathways: the depletion of *xpf-1* further decreased the fusion phenotype in *polq-1,cku-70;rad-51;him-5* (Figure 4E-G; P=0.0266).

### Regulation of repair pathways in *rad-51;him-5*

Having ascertained that c-NHEJ, TMEJ, and SSA can be active in the meiotic germ line, at least when HR is impaired, we next wanted to ask whether these pathways contribute to repair in the *rad-51;him-5* mutant background. None of the mutations that impair c-NHEJ, TMEJ or SSA by themselves or in pairwise combinations altered the number and morphology of DAPI bodies in *rad-51;him-5* (Figure S5 A-E, G; Kruskal-Wallis, *P*>0.43). However, a *rad-51;him-5* strain with mutations in all three pathways had statistically significantly fewer DAPI-bodies per diakinesis oocyte (Figure S5 F, G; Kruskal-Wallis test, P=0.0055). These results reveals that the repair pathways act redundantly to repair DSBs in the absence of both RAD-51 and HIM-5. It also confirms that DSBs are induced in the *rad-51;him-5* mutant germ cells (see discussion).

The intact appearance of diakinesis-stage chromosomes in *rad-51;him-5*, in particular the lack of chromosome fragments, could also be attributed to the low level of DSBs that result from loss of *him-5* function. We therefore wanted to reexamine the triple, quadruple, and quintuple mutant lines after irradiation, thereby compensating for the lack of DSBs. As discussed above, exposure of *rad-51;him-5* mutant animals to 10 Gy IR led to a statistically significant decrease in the number of DAPI bodies in diakinesis oocytes, with the frequent appearance of one or more chromosome fusions and occasional small DNA fragments (Figure 2D). We note that this dose of IR did not dramatically alter the appearance of *rad-51* diakinesis oocytes (Two-tailed Mann-Whitney, *P*>0.49), suggesting differences in the behavior of chromosomes in *rad-51* versus *rad-51;him-5*.

While the depletion of c-NHEJ eliminated some of the fusions that were formed in irradiated *rad-51;him-5* (10.56 DAPI-bodies on average in *cku-80;rad-51;him-5* vs. 9.87 in *rad-51;him-* 5) (Figure 5 A, G; Kruskal-Wallis, *P*=0.0256), the depletion of TMEJ or SSA, alone or in combination, did not have a significant effect on nuclear DAPI-body content (Figure 5B, C, G; Kruskal-Wallis, *P*>0.05), suggesting that these two pathways were not essential to the repair of IR-generated breaks in the *rad-51;him-5* background. In contrast, the presence of functional c-NHEJ was key; any mutant combination that lacked a component of the Ku-complex showed a significant difference in DAPI-body counts compared to *rad-51;him-5* (Figure 5D-G; Kruskal Wallis, *P*<0.03). We note that in *cku-80;rad-51;him-5*, loss of c-NHEJ function prevented the formation of chromosomal fusions upon IR exposure, as observed by an increase in the average number in DAPI-bodies compared to *rad-51;him-5*. By contrast, co-depletion of c-NHEJ and TMEJ and/or SSA led to a decrease in DAPI-body numbers (reflecting an increase in chromosome fusions; Figure 5D, F, G). This suggests that while c-NHEJ, TMEJ and SSA pathways all act as backups to repair DSBs in *rad-51* mutants, their use is disturbed by the additional loss of *him-5*.

**Figure 5.**
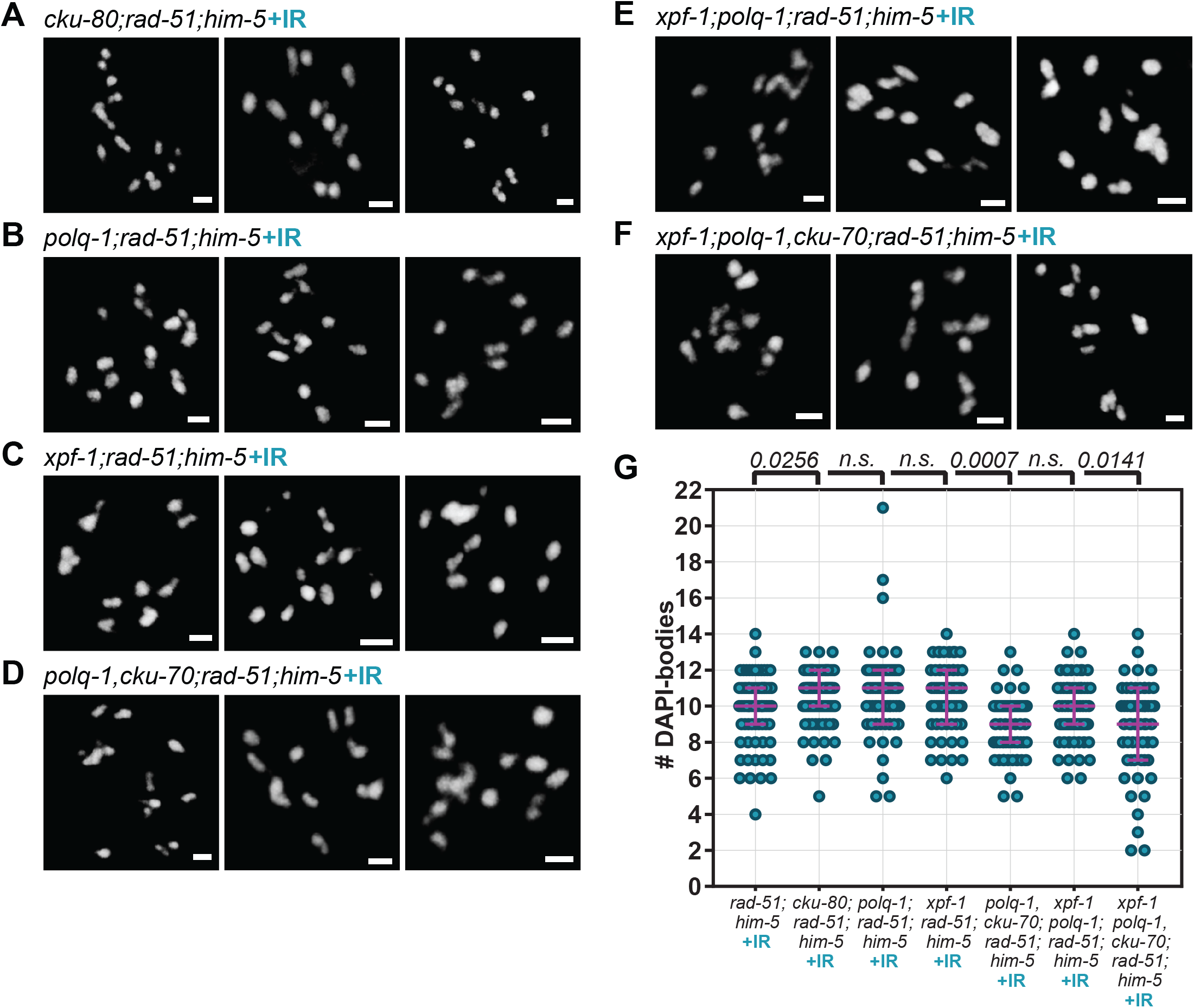
c-NHEJ is the major repair pathway for IR-induced breaks in *rad-51;him-5* mutants. (A-F) Representative images and (G) Quantification of DAPI-stained diakinesis nuclei of two-day-old, irradiated (+IR) adults. Scale bars= 2*μ*m. (G) Magenta bars indicate the median and the interquartile range. Kruskal-Wallis multiple comparisons to *rad-51;him-5* (n.s.: not significant, P>0.05). *rad-51(lg8701);him-5(e1490*) (n=71), *cku-80(ok851);rad-51(lg8701);him-5(e1490*) (n=80), *polq-1(tm2027);rad-51(lg8701);him-5(e1490*) (n=67), *xpf-1(e1487);rad-51(lg8701);him-5(e1490*) (n=61), *polq-1(tm2026),cku-70(tm1524);rad-51(lg8701);him-5(e1490*) (n=63), *xpf-1(e1487);polq-1(tm2026);rad-51(lg8701);him-5(e1490*) (n=64), *xpf-1(e1487);polq-1(tm2026),cku-70(tm1524);rad-51(lg8701);him-5(e1490*) (n=59).

### Exploring the accessibility of alternative repair machineries to DSB ends

The dependence of TMEJ and SSA, but not c-NHEJ, on the presence of *him-5* led us to explore the differences in the requirements for these pathways. Specifically, we honed in on the role of resection in repair pathway choice. TMEJ and SSA rely on stretches of DNA sequence homology which are typically found between repetitive elements in the genome (Sugawara *et al*. 2000). The availability of such stretches of ssDNA to the repair machinery requires 5’ to 3’ resection of DSB ends. Upon DSB formation, the activity of MRN/COM-1 generates short resected ends through the clipping of SPO-11 oligos. In mitotic cells, extensive resection requires the action of the exonuclease EXO-1 (Lemmens and Tijsterman 2011), which has also been implicated in meiotic repair in nematodes and other species (Mimitou and Symington 2008; Zhu *et al*. 2008; Nimonkar *et al*. 2011; Lemmens *et al*. 2013; Yin and Smolikove 2013). c-NHEJ, by contrast, preferentially uses unresected ends. We thus hypothesized that the modulation of TMEJ and SSA by *him-5* could be a consequence of a role for *him-5* in EXO-1-mediated resection. To test this hypothesis, we compared the impact of depleting *exo-1* or *mre-11* in *rad-51* and *rad-51;him-5* mutant animals.

To our knowledge, the effect of depleting short and long resection specifically in a *rad-51* mutant background has not been assayed. Consequently, we first examined the impact of loss *of exo-1 or mre-11* in *rad-51* mutant animals. The *mre-11S* mutant animals exhibit chromosome fusion phenotypes similar to *rad-51* mutant animals, yet were more likely to result in >6 DAPI bodies (Figure 6C, Figure S6 D, F). With the addition of 10Gy IR, *mre-11S* gave a more severe fusion phenotype than *rad-51* (Figure 6A-C). The spectrum of fusion phenotypes conferred by these mutations likely reflects differences in repair pathway usage in each mutant. As previously reported, loss of *exo-1* function did not affect meiotic crossover formation (Figure S6 A-C), nor did it alter the phenotype of *mre-11* (Figure S6 D-F; (Yin and Smolikove 2013)). The nuclear content in DAPI-stained bodies in *rad-51;mre-11S* (5.90 on average) and in *exo-1;rad-51* (5.21) were both significantly higher than in *rad-51* single mutants (4.49; Kruskal-Wallis, *P*<0.012). The partial suppression of the *rad-51* fusions suggests that both short and long resection contribute to fusions that form in *rad-51* mutant animals (Figure 6D-F).

**Figure 6.**
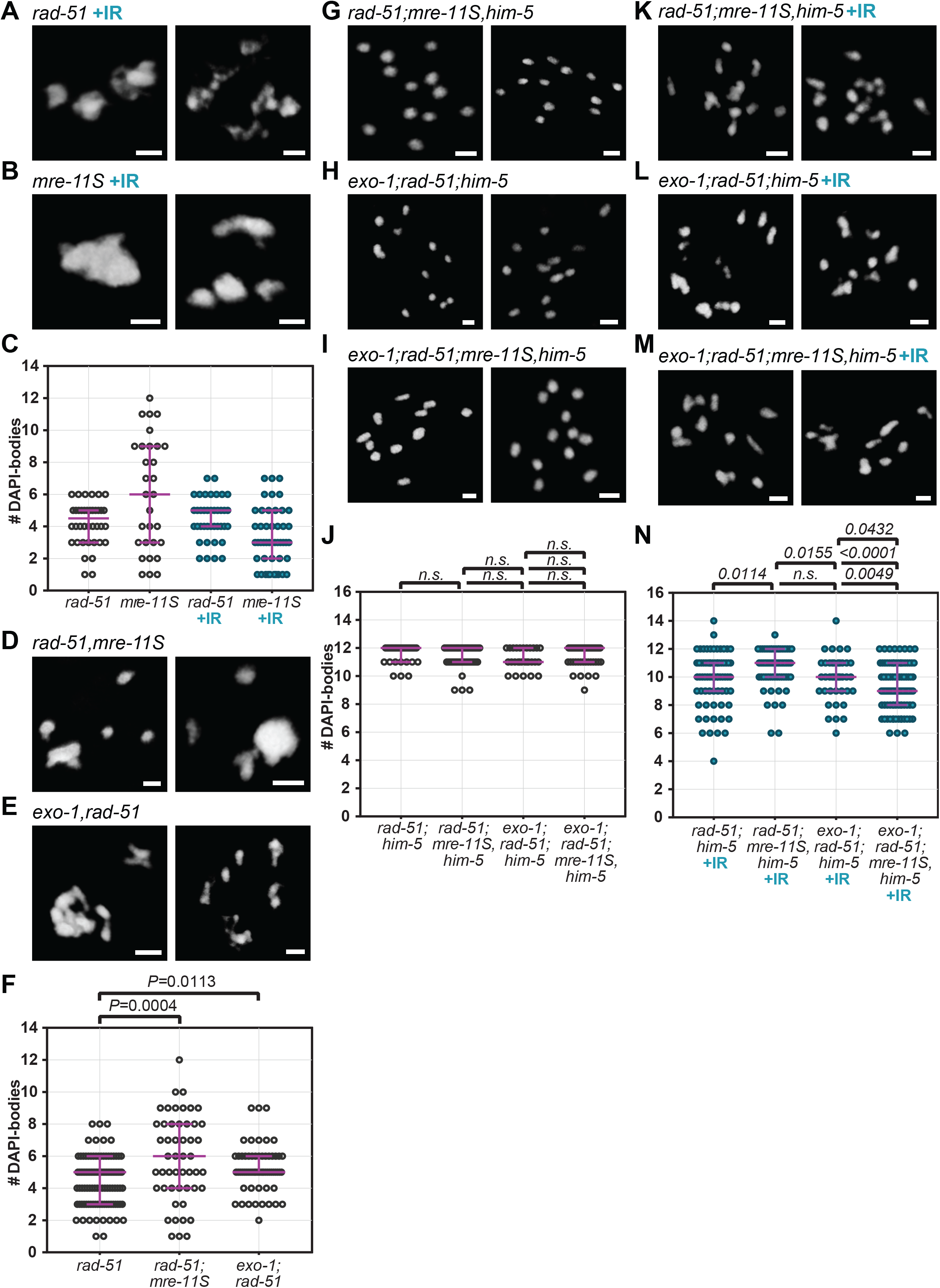
DSB repair is regulated differently in *rad-51* and *mre-11S* mutants. (A-C) IR-induced breaks do not change the phenotype of *rad-51* mutants, while they do affect *mre-11S* mutants. Shown are DAPI-bodies in representative diakinesis nuclei of unirradiated and irradiated (+IR) animals of indicated genotypes. (D-F) Depleting *mre-11S* or *exo-1* in *rad-51* mutants affects chromatin morphology, suggesting alterations in how DSBs are repaired. (A, B, D, E) Images of diakinesis nuclei of two-day-old adults of indicated genotypes. (C, F) Quantification of DAPI-stained bodies in diakinesis nuclei. (G-N) *him-5* does not influence resection, but MRE-11 and EXO-1 act in concert for the repair of IR-induced breaks. (G-J) *mre-11S* or *exo-1* mutations have no effect on diakinesis chromosome morphology in the *rad-51;him-5* background. (K-N) By contrast, *mre-11S* and *exo-1* affect repair of IR-induced damage, independently of *him-5*. (G-I, K-M) Pictures show DAPI-stained diakinesis nuclei of adults on day 2 of adulthood. Scale bars= 2*μ*m. (J, K) Quantification of DAPI-bodies in diakinesis nuclei of indicated genotypes. Magenta bars indicate median and interquartile range. P-values calculated from Kruskal Wallis tests (*n.s*.: not significant, *P*> 0.15). *rad-51(lg8701*) (n=40), *mre-11S(iow1*) (n=30), *rad-51(lg8701)+IR* (n=39), *mre-11S(iow1)+IR* (n=47). F: *rad-51(lg8701*) (n=106), *rad-51(lg8701);mre-11S(iow1*) (n=50), *exo-1(tm1842);rad-51(lg8701*) (n=63). J: *rad-51(lg8701);him-5(e1490*) (n=33), *rad-51(lg8701);mre-11S(iow1),him-5(e1490*) (n=57), *exo-1(tm1842);rad-51 (lg8701);him-5(e1490*) (n=28), *exo-1(tm1842);rad-51(lg8701);mre-11S(iow1),him-5(e1490*) (n=50). N: *rad-51(lg8701);him-5(e1490)+IR* (n=71), *rad-51(lg8701);mre-11S(iow1),him-5(e1490)+IR* (n=60), *exo-1(tm1842);rad-51 (lg8701);him-5(e1490)+IR* (n=39), *exo-1(tm1842);rad-51(lg8701);mre-11S(iow1), him −5(e1490)+I* R (n=101).

In the *rad-51;him-5* double mutant context, the depletion or co-depletion of *mre-11S* and *exo-1* did not affect the univalent chromosomes (Figure 6G-J). Upon IR however, the depletion of *mre-11S* appears to cause a significant decrease in the formation of fusions (10.63 DAPI-bodies on average in *rad-51;mre-11S,him-5* vs 9.87 in *rad-51;him-5*, Kruskal-Wallis test, *P*=0.0114), while the depletion of *exo-1* alone did not have a significant effect (9.90 bodies per nucleus, *P*>0.78) (Figure 6K-N). Interestingly however, the co-depletion of *mre-11S* and *exo-1* caused significantly more fusions to occur (9.24 bodies per nucleus on average) compared to *rad-51;him-5* double mutants, but also compared to *rad-51;mre-11S,him-5* and *exo-1;rad-51:him-5* triples (*P*<0.05, Figure 6G, H). While this data does not allow us to draw any conclusion about a role of *him-5* on resection, we note a concerted action of MRE-11 and EXO-1 on dirty breaks. This is confirmed by the observation that, as discussed above, while the depletion of *exo-1* did not alter the phenotype of *mre-11S* mutants in non-IR conditions, it did have an effect upon IR (Figure S6 F). Indeed, fewer fusions were formed in irradiated *exo-1;mre-11S* compared to irradiated *mre-11S* (4.91 vs. 3.32 DAPI-bodies per nucleus on average, Mann-Whitney test, *P*=0.0050). This emphasizes the fact that the nature of the DSB influences the way it will be repaired.

### The DSB-promoting machinery regulates downstream repair events

Having established a role for HIM-5 in influencing repair pathway choice, we next wanted to determine if this function was specific to HIM-5 or whether this is a more general feature of DSB-promoting factors. Mutations in *dsb-2* confer a partial defect in DSB formation that worsens with maternal age, causing a shortage in chiasmata and increased nondisjunction (Rosu *et al*. 2013). In accordance with previous reports, we observed an average of 9.33 DAPI-bodies per nucleus in *dsb-2* mutant (day 2 post-L4; Figure 7A). Upon treatment of *dsb-2* mutant worms with 10 Gy IR, the average number of DAPI-bodies was restored to 5.93 intact ovoid structures, a level which is indistinguishable from wild-type worms (Figure 7A, B; Mann-Whitney test, *P*=0.5567). We discovered that diakinesis nuclei in *dsb-2;rad-51* double mutants contained ~12 DAPI-bodies (Figure 7C, D). This phenotype is similar to what is observed in *rad-51;him-5*. Further, IR of *dsb-2;rad-51* failed to recapitulate the *rad-51* mutant phenotype and was instead not significantly different from unirradiated *dsb-2;rad-51* (Figure 7C, E; *P*=0.1674). Moreover, we observed that SPO-11-mediated DSBs are still formed in *dsb-2;rad-51*. Indeed, 70% of *dsb-2;rad-51,rec-8* triple mutant diakinesis nuclei contained fragments of chromatin (n=23), contrasting with the absence of fragments in *spo-11,rec-8* nuclei (n=44), where DSBs are abolished (Figure 7F, G). We were thus able to observe the same suppression of chromosome fusions that occur in *rad-51* mutants as in *rad-51;him-5* by depleting *dsb-2*. Again, we show that this is not due to an abrogation of DSBs.

**Figure 7.**
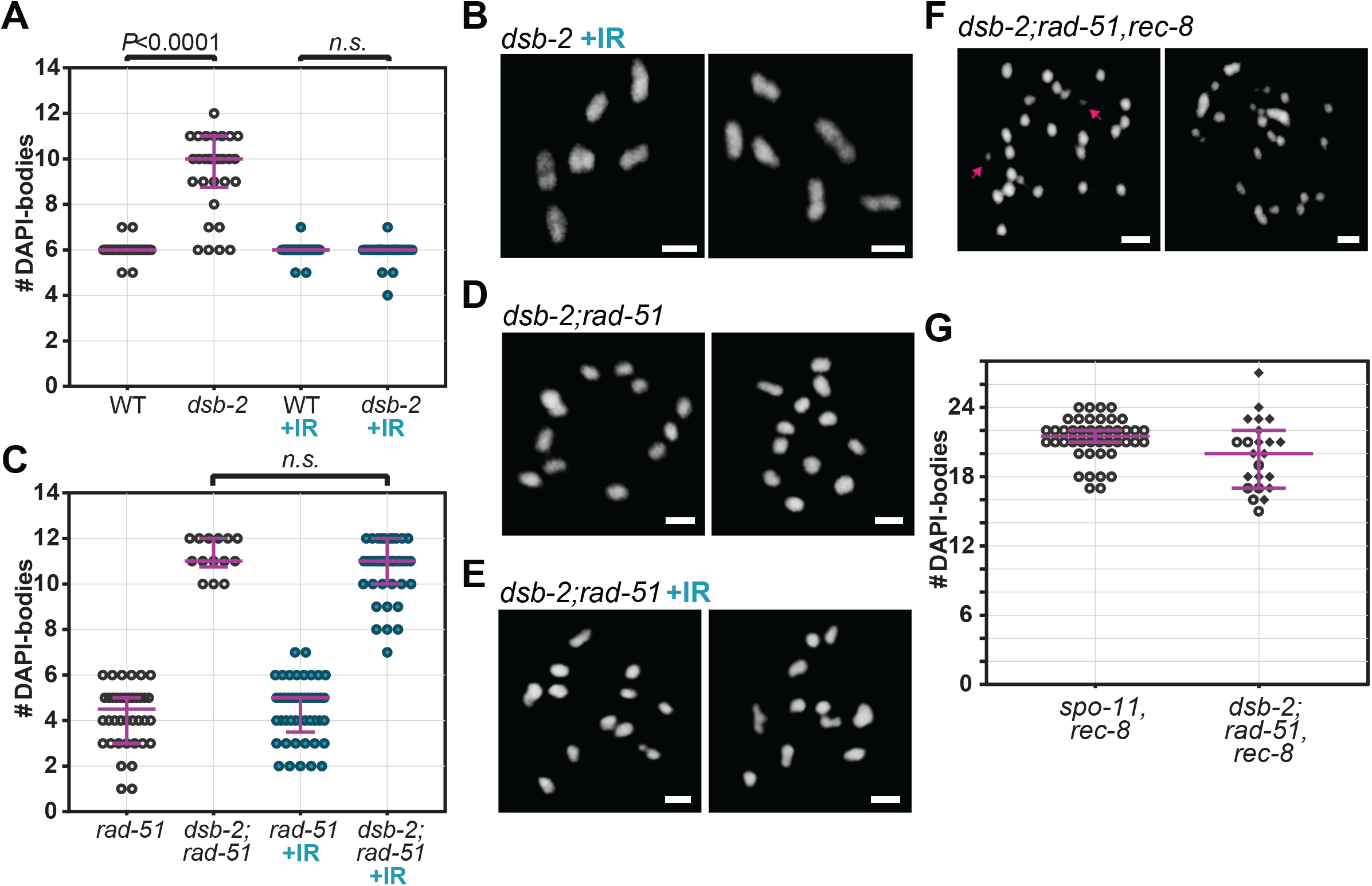
Loss of *dsb-2*function impacts repair in *rad-51* mutants, similar to *him-5*. (A, B) Control showing that IR suppresses the univalent phenotype induced by loss of *dsb-2* (A) Quantification of diakinesis oocytes of indicated genotypes. Wild type (WT, n=45), *dsb-2(me96*) (n=30), WT+IR (n=52), *dsb-2(me96*) +IR (n=44). Two-tailed Mann-Whitney tests are indicated on top of the graph as exact P-values (n.s.: not significant, P=0.5567). (B) Representative images of diakinesis oocytes in irradiated two-day-old, *dsb-2* mutant animals. (C-E) Depletion of *dsb-2* suppresses fusions in *rad-51* mutants. (C-E) Diakinesis nuclei in *dsb-2;rad-51* double mutants contain 12 intact univalents that are largely unaffected by IR-induced breaks. (C) Quantification and (D, E) Representative images of DAPI-stained diakinesis nuclei. *rad-51(lg8701*) (n=40), *dsb-2(me96);rad-51(lg8701*) (n=14), *rad-51* +IR (n=45), *dsb-2;rad-51(lg8701*) +IR (n=39). two-tailed Mann-Whitney test (n.s.: not significant, P>0.16). (F, G) SPO-11-mediated breaks are formed in *dsb-2;rad-51* as seen by (F) chromatin fragments in the nuclei of *dsb-2;rad-51,rec-8* (compare *spo-11,rec8* in Figure 3A). (G) Quantification of DAPI bodies in diakinesis oocytes of *spo-11(me44),rec-8(ok978*) (n=44), *dsb-2(me96);rad-51(lg8701),rec-8(ok978*) (n=23). Diamonds correspond to nuclei containing at least one fragment of chromatin. Scale bars in images = 2*μ*m. Magenta bars in dot plots indicate median and interquartile range.

## DISCUSSION

### Multiple error-prone DSB repair pathways are active in the meiotic germ line

Throughout the cell cycle, different DSB repair pathways are utilized to ensure the accurate repair of genetic information. In mitotically cycling cells, c-NHEJ is the dominant repair pathway in G1; whereas HR is preferentially utilized in S and G2 when a sister chromatid template is available. In meiotic cells, HR appears to be the major repair pathway since it is the only repair pathway that is considered “error-free” and can ensure the integrity of the genome across generations. It is well documented that c-NHEJ can step in to repair DSBs when HR is impaired during meioisis (Nairz and Klein 1997; Stark *et al*. 2004; Smolikov *et al*. 2007b; Lemmens *et al*. 2013; Yin and Smolikove 2013). The contributions of other DSB repair pathways are less well defined. Our data reveal multiple redundancies between c-NHEJ, TMEJ, and SSA during DSB repair in the meiotic germ line. To our knowledge, this is the first evidence that all of these pathways are active in meiotic germ cells and contribute as a failsafe mechanism to protect genomic integrity.

Checchi *et al*. revealed that in the absence of HR, SSA can repair DSBs on the male X chromosome in *C. elegans*. We provide additional data to support a role for SSA as an alternative repair pathway showing that SSA contributes to repair in the hermaphrodite oocyte germ lineage. Depletion of *xpf-1* had the greatest impact on diakinesis DNA morphology in the *rad-51* mutant background, suggesting that SSA is the major default pathway of repair in the absence of RAD-51 and strand invasion functions. This may simply be a consequence of having long-resected ends which would normally promote strand invasion. c-NHEJ and TMEJ play important, but more minor roles in repair when *rad-51* function is impaired, consistent with their preference for shorter or blunt DNA ends.

The redundancy between TMEJ, SSA, and c-NHEJ in meiotic DSB repair in the absence of HR function suggests that the traditional dichotomy between c-NHEJ and HR repair is more complex. Our data suggest that a key decision point is between use of the ssDNA filament in SSA or HR. How is the bias for HR accomplished? Since SSA requires homologous direct repeats, it may be disfavored during meiosis in which programmed DSBs are directed away from repetitive genomic sequences (Sasaki *et al*. 2010). Alternatively, specific components of the meiotic break and processing machinery may directly inhibit SSA. Another possibility is that protein(s) required for SSA are not expressed or active during the leptotene to early pachytene stages of meiosis when crossovers are established.

### HIM-5 and Pathways Choice Functions

In the absence of HIM-5 proteins, repair pathway bias in *rad-51* mutant animals shifted. c-NHEJ now became important for repair; whereas TMEJ and SSA adopt more auxiliary functions. Thus, HIM-5 function appears to promote use of TMEJ and SSA, at least when strand exchange is impaired. Several recent studies have pointed out that coupling DSB formation to downstream resection and repair facilitates repair through HR (Lemmens *et al*. 2013; Mateo *et al*. 2016; Girard *et al*. 2018). We previously showed that HIM-5, redundantly with the p53 homolog CEP-1, prevents access to c-NHEJ, a function also accomplished by COM-1/CtIP and MRN. We now show that HIM-5 also promotes homology-directed repair, either through HR or TMEJ/SSA.

We consider several different models for how HIM-5 can influence repair pathway choice. First, since HIM-5 is a chromatin-associated protein, it could regulate transcription or chromatin recruitment of key factors that promote HR and TMEJ/SSA. Decreased expression of these factors in *him-5* mutant animals would impair HDR pathways and shift repair capacity to the more error-prone c-NHEJ. Second, HIM-5 could directly inhibit components of the c-NHEJ pathway, redundantly with CEP-1, under wild-type conditions. Third, based on our analysis of *exo-1* and *mre-11*, we envision a role for HIM-5 in modulating the resection machinery. MRE-11, like HIM-5, functions in both DSB formation and resection. By contrast, EXO-1 is dispensable for both under wild-type situations. In *rad-51;him-5*, however, EXO-1 function became important for DNA repair in an MRE-11-dependent fashion. The shift in importance for EXO-1 could reflect a function for HIM-5 in coordinating assembly of the DSB machinery, including MRE-11. Consistent with this model, our data suggest that DSB-2, like HIM-5, may be involved in the promotion of HR-mediated DSB repair. Although deeper analysis of multi-mutants will be necessary to know if DSB-2 influences the same events as HIM-5, these results raise the possibility that the DSB-promoting machinery directly couples DSB induction with resection and also with the downstream events that promote HR during meiosis.

### Timing of DSB break repair in meiosis shifts during early to late pachytene

A possibility that is not incompatible with other models is that HIM-5 acts as a timer for meiotic events, ensuring the appropriate formation of meiotic DSB formation in leptotene/zygotene and subsequent HR repair and crossover commitment in early pachytene. In this scenario, the loss of HIM-5 functions would delay DSB induction and/or repair until mid-late pachytene, leading to both a decrease in CO number (due a shorter window of opportunity for DSB formation) and the potential of aberrant repair by alternative DSB pathways.

This model is consistent with our prior data that loss of *him-5* function delayed the appearance, accumulation, and total number of RAD-51 foci (Meneely *et al*. 2012; Mateo *et al*. 2016) and delayed the maturation of COs (Machovina *et al*. 2016). The timing model could also explain the shift in the dependence for *exo-1* function. Smolikove and colleagues (Yin and Smolikove 2013) showed that *exo-1* can contribute to CO repair when both *mre-11* and c-NHEJ are mutated but that accumulation of RAD-51 in these animals is delayed until late pachytene, suggesting EXO-1 resection is either slower than MRE-11-induced resection or that EXO-1 is not active until late pachytene (LP). The contribution of *exo-1* to repair in *rad-51;him-5* might therefore be explained by either delayed timing of DSB formation or delayed end processing in *him-5*. Early pachytene may be conducive to DSB formation and repair via HR, whereas later events may be channeled to alternative repair pathways.

Further support for differences in DNA repair pathway accessibility throughout prophase comes from the observation that RAD-51 foci persist in *rad-54* mutants in LP nuclei (Checchi *et al*. 2014), well after RAD-51 foci have disappeared in wild type ((Koury *et al*. 2018) and Li and Yanowitz, unpublished data). If the alternative repair pathways were active early in pachytene, RAD-51 should have disappeared with near normal kinetics. The late removal of RAD-51 suggests that these pathways are not active until late pachytene. Coincidently, the mid-to-late pachytene (MP-LP) transition marks a change in HR repair from *rad-50-* dependent to *rad-50* independent, which is thought to correspond to a shift from the interhomolog, CO pathway to an inter-sister, error-free HR repair pathway (Hayashi *et al*. 2007). The MP-LP transition is also associated with the loss of DSB competency, assuring that additional SPO-11 mediated meiotic breaks are not made. Given that both HIM-5 and DSB-2 are components of the DSB machinery and their accumulation in the nucleus is lost in LP, we suggest that barriers to alternative repair are also lifted in LP, perhaps as a consequence of the loss of the DSB proteins. The exact coordination of these events in the germ line will be an area of future investigation.

### Evidence for additional back-up repair pathways

Surprisingly, *xpf-1;polq-1;cku-70;rad-51* and *xpf-1;polq-1;cku-70;rad-51;him-5* animals that are impaired for SSA, TMEJ, c-NHEJ, and HR functions exhibit genomes that are surprisingly intact. One might have anticipated seeing twelve large chromosome fragments corresponding to each of the unattached homologs and a large fraction of small to medium sized fragments corresponding to the pieces of chromosomes end that are cut by SPO-11. However, even the addition of IR in the *him-5* background leaves a fairly intact genome. One possible explanation for these results is that we have only partially impaired the function of one or more of these DSB repair pathways, despite the use of null alleles. Alternatively, the mild phenotype may result from the activity of additional pathways that ligate chromosomes, such as other microhomology-mediated mechanisms. We note that even in the presence of additional DSBs, *xpf-1;polq-1;cku-70;rad-51* and *xpf-1;polq-1;cku-70;rad-51;him-5* mutant germ cells are not identical, suggesting that HIM-5 function still biases repair outcomes even when SSA, TMEJ, NHEJ, and HR are not functional.

## DIRTY vs CLEAN DSBs

Examination of our data with and without IR, clearly points to distinct methods of repair for SPO-11-induced vs IR-induced lesions. A clear example of this is seen in the comparison of *rad-51* and *mre-11S* in the presence or absence of IR. Whereas IR has little impact on *rad-51* (compare Figures 1D versus Figure 2C), IR both reduced the variability of diakinesis phenotypes in *mre-11S* and led to more massive chromosome clumps compared to *rad-51*. The differences in the *mre-11S* background can be explained if the covalent attachment of SPO-11 to the DSBs occludes binding to Ku proteins until endonucleolytic cleavage. This would assist to bias repair towards HDR pathways. The damage created by IR tends to be staggered DSBs breaks that are ideal substrates for TMEJ or that would be rapidly processed into blunt ends for Ku complex recruitment for c-NHEJ.

Another example of the difference in SPO-11 vs. IR breaks comes from examination of the *xpf-1* mutation in *rad-51* and *rad-51;him-5* mutant backgrounds. In *rad-51* mutant animals, *xpf-1* mutation confers the greatest suppression of fusion phenotypes. In *rad-51;him-5*, loss of *xpf-1* has no effects with or without IR and, perhaps more significantly, loss of *xpf-1* does not enhance the suppression conferred by *polq-1;cku-70* double mutants both with and without IR. The differences in the processing of SPO-11 and IR-induced DSBs serves as a cautionary tale for a field that relies on IR as a surrogate for DSBs since they can be converted to COs.

## ACKNOWLEDGMENTS

The authors wish to thank Logan Russell for careful reading of the manuscript and members past and present of the Yanowitz lab for intellectual insights into the project. The work was funded by NIH grant: R01GM104007 to J.L.Y.; N.M. was also funded in part by the Magee-Womens Auxiliary Bright Star Award. Some strains were provided by the CGC, which is funded by NIH Office of Research Infrastructure Programs (P40 OD010440).

---

**Figure S1.**
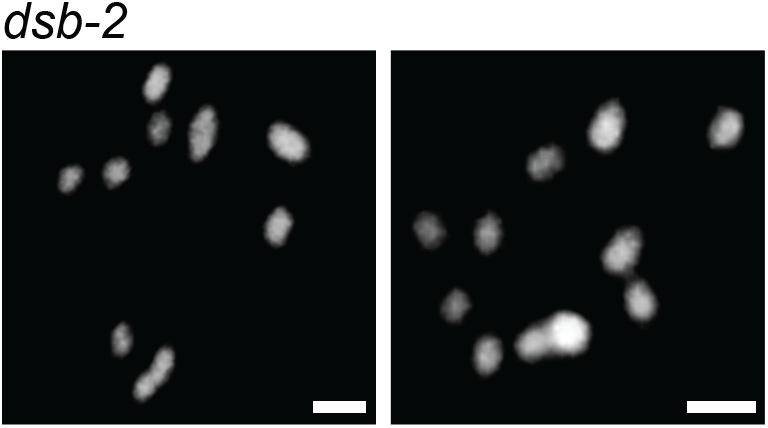
The *dsb-2* mutant shows a shortage in chiasmata. The lack of chiasmata revealed by univalents in diakinesis nuclei, as previously reported (Rosu et al. 2013). Representative DAPI-stained diakinesis nuclei of *dsb-2* mutants at day 2 of adulthood. Scale bars= 2*μ*m.

**Figure S2.**
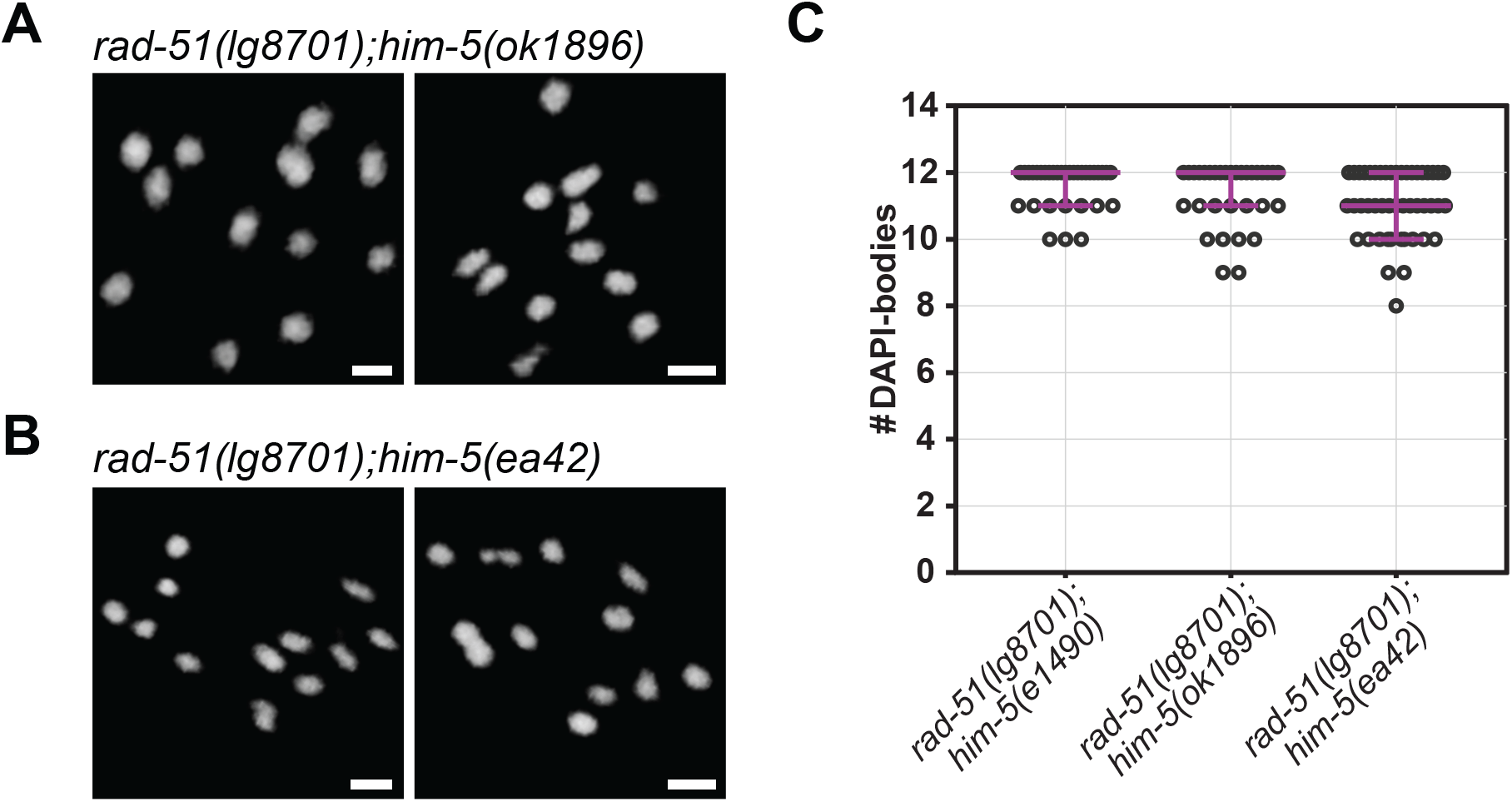
Suppression of *rad-51* chromosomes fusions by *him-5* is not allele-specific. (A-C) Multiple alleles of *him-5* suppress the formation of chromosomal fusions that normally arise when *rad-51* is mutated. (A, B) Represenative images and (C) Quantificationof DAPI-stained diakinesis nuclei in two-day-old adults of indicated genotypes. Scale bars= 2*μ*m. Magenta bars indicate the median and the interquartile range. P-values from Kruskal-Wallis multiple comparison are indicated on top of the graph (n.s.: not significant, P>0.11). *rad-51(lg8701);him-5(e1490*) (n=33), *rad-51(lg8701);him-5(ok1896*) (n=32) and *rad-51(lg8701);him-5(ea42*) (n=47).

**Figure S3.**
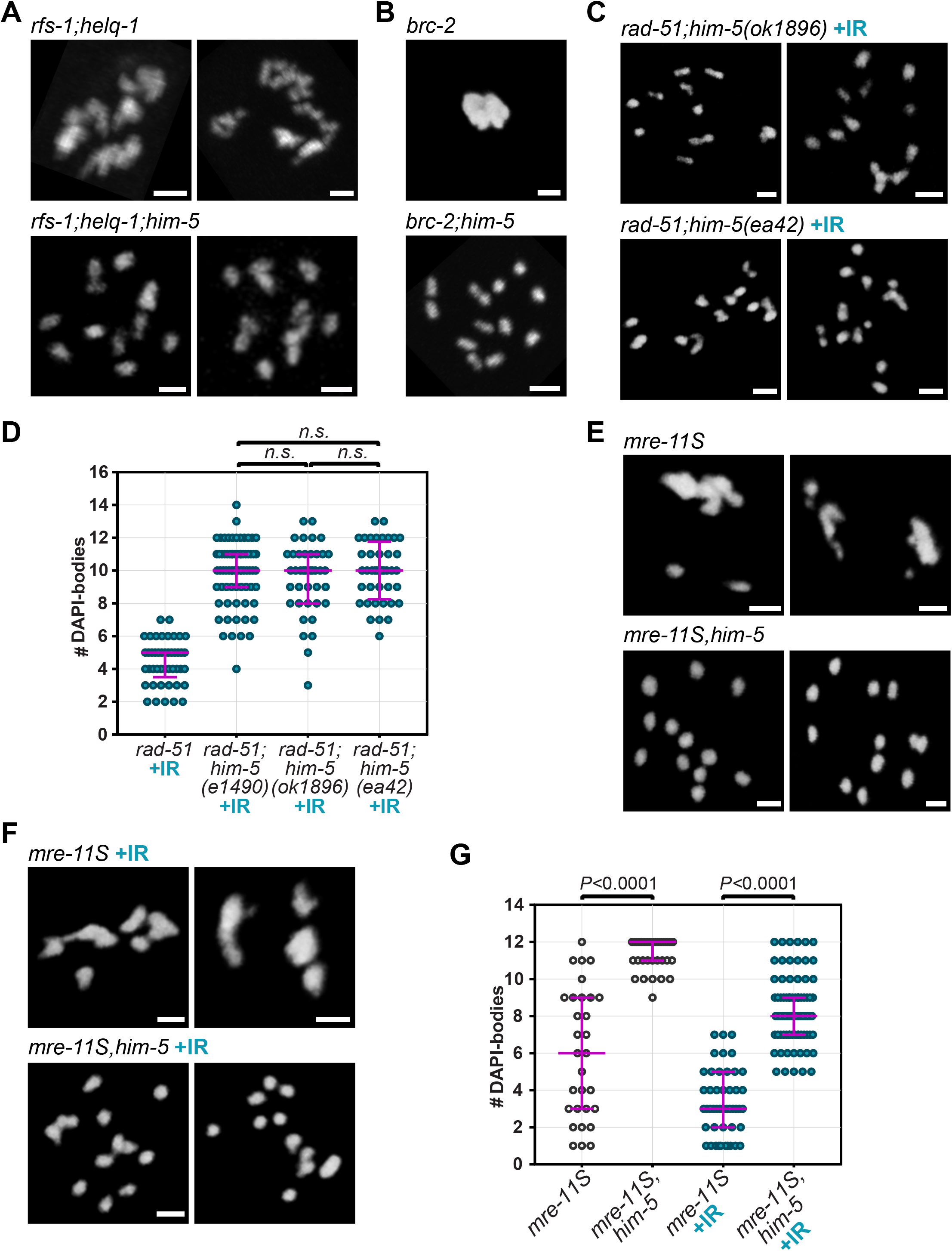
The depletion of *him-5* suppresses chromosomes fusions caused by mutations in genes encoding proteins required for strand-exchange. (A) *rfs-1;helq-1* and (B) *brc-2* mutant animals exhibit chromosomal clumping at diakinesis that is suppressed by the depletion of *him-5*. (C, D) Irradiation of *rad-51;him-5* does not restore a *rad-51-like* phenotype (C) Representative images and (D) Quantification of diakinesis nuclei from two-day-old irradiated animals: *rad-51(lg8701*) (n=45), *rad-51(lg8701);him-5(e1490*) (n=71), *rad-51(lg8701);him-5(ok1896*) (n=40), and *rad-51(lg8701);him-5(ea42*) (n=40). Kruskal-Wallis multiple comparison indicate that irradiated double mutants are indistinguishable from each other (n.s.: not significant, *P*>0.9999). (E-G) Diakinesis nuclei of *mre-11S,him-5* double mutants contain univalents than are not suppressed by IR-induced breaks. (E-F) Diakinesis nuclei of two-day-old adults without and with (+) IR. Scale bars= 2*μ*m. (G) Quantification of DAPI bodies in diakinesis nuclei of different genotypes: unirradiated (clear dots) *mre-11S(iow1*) (n=30), *mre-11S(iow1);him-5(e1490*) (n=39), and irradiated (+IR, blue dots) *mre-11S(iow1*) (n=47) and *mre-11S(iow1);him-5(e1490*) (n=79). P-values calculated by two-tailed Mann-Whitney tests. Magenta bars indicate median and interquartile range (D, G).

**Figure S4.**
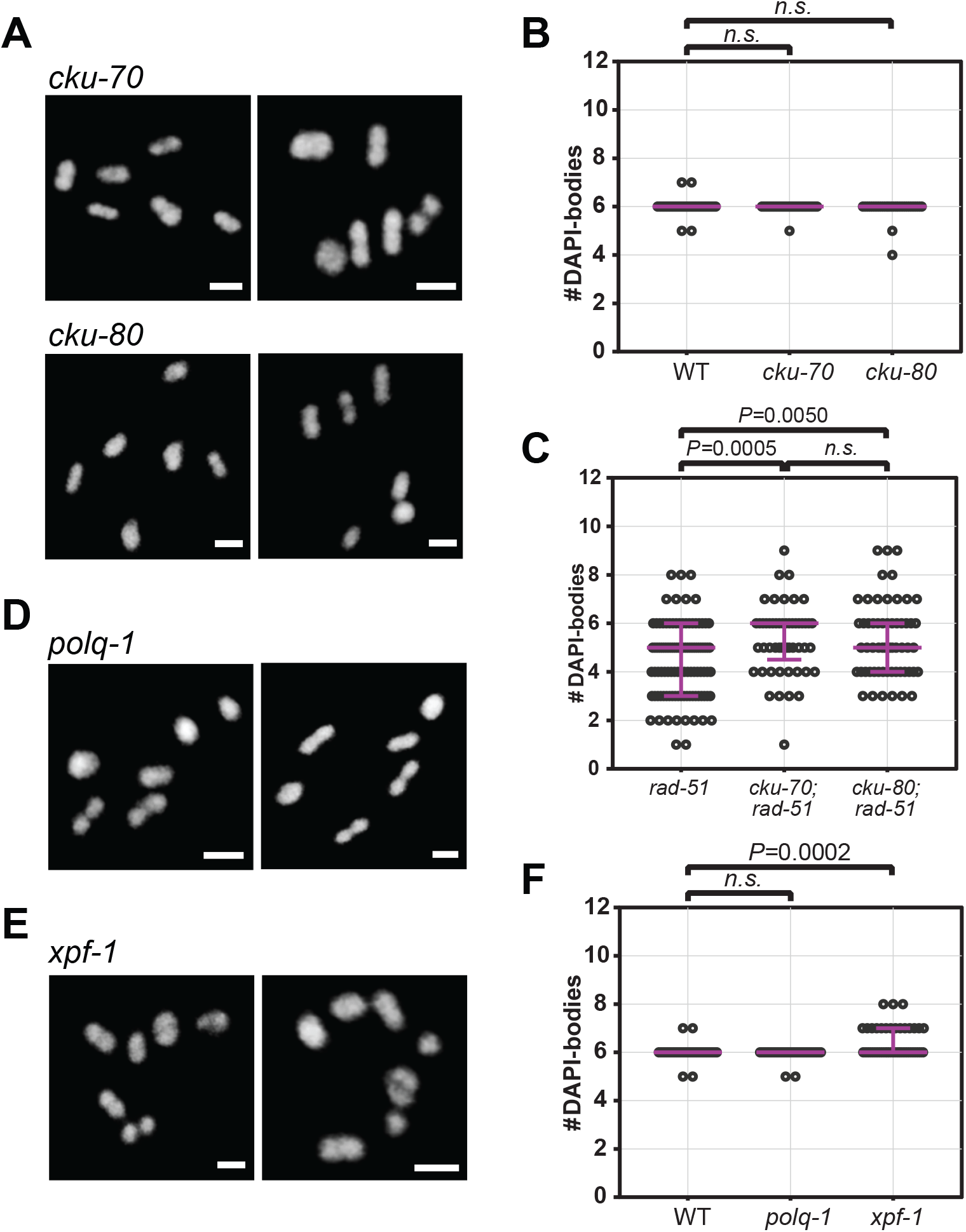
Mild to no crossover defects are observed in animals carrying single mutations in non-HR repair genes. (A, B) Diakinesis nuclei of mutants for Ku-complex components contain six ovoid DAPI-stained bodies and are indistinguishable from nuclei of wild-type worms. (A) Representative images and (B) Quantification of diakinesis oocytes in one-day-old adults. (C) Impairment of c-NHEJ partially suppresses the formation of chromatin fusions in *rad-51* mutant animals, increasing the number of DAPI-dense structures in diakinesis nuclei. (D) Depletion of *polq-1* does not affect crossover formation by itself as six bivalents are seen in all nuclei. (E) Univalents are seen occasionally in xpf-1 mutant animals. DAPI-stained diakinesis nuclei at day 2 of adulthood. Scale bars= 2*μ*m. (F) Quantification of DAPI-bodies in diakinesis nuclei. Magenta bars indicate the median and the interquartile range. Two-tailed Mann-Whitney tests (n.s.: not significant, P>0.19). Wild type (WT, n=45), cku-70(tm1524) (n=40), cku-80(ok851) (n=32), polq-1(tm2026) (n=45), xpf-1(e1487) (n=55), rad-51(lg8701) (n=106), cku-70(tm1524);rad-51(lg8701) (n=53), cku-80(ok851);rad-51 (l g8701) (n=58).

**Figure S5.**
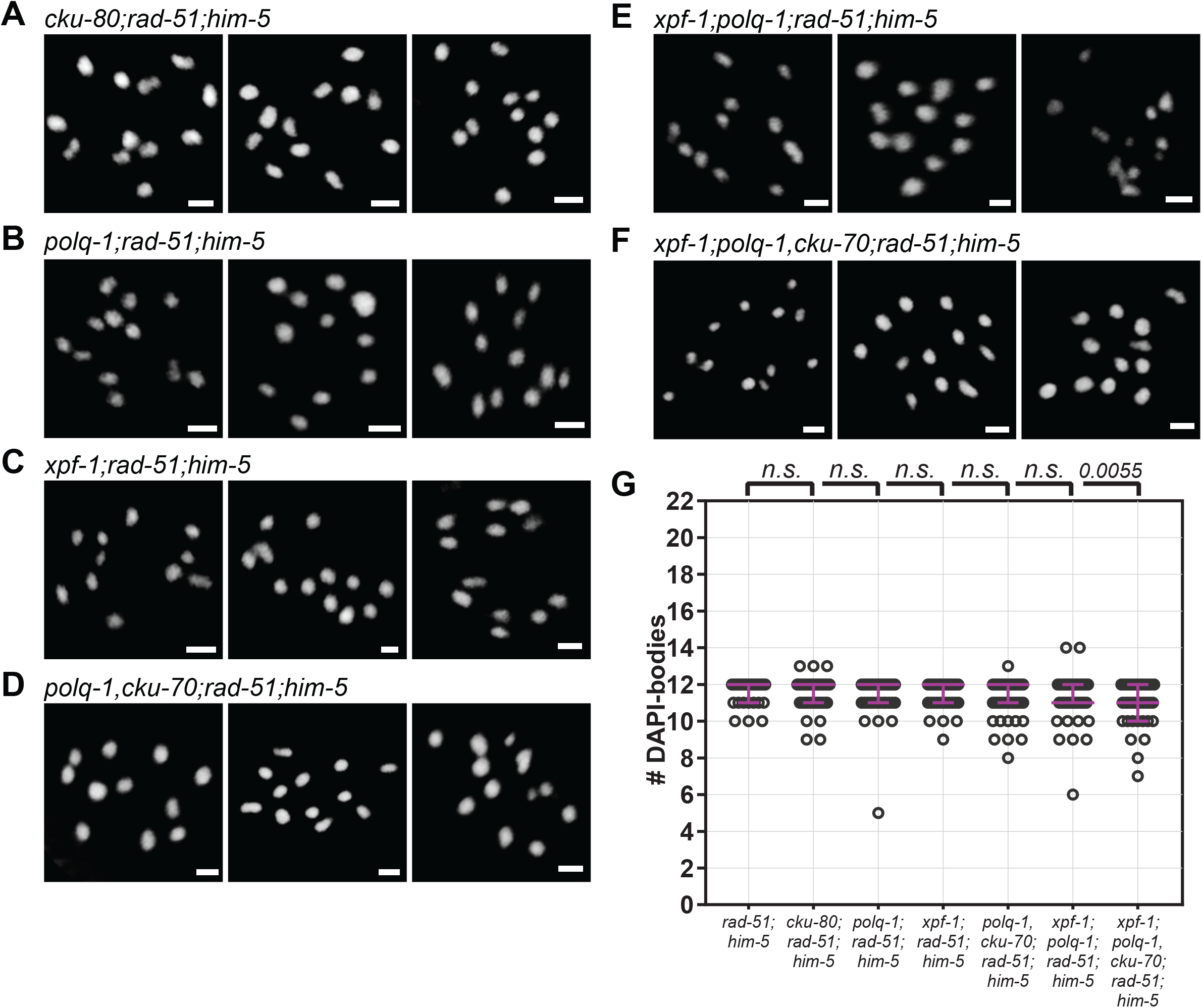
c-NHEJ, TMEJ, and SSA function redundantly to control repair in *rad-51;him-5* mutants under low DSB conditions (A-F) DAPI-stained diakinesis nuclei of two-day-old, irradiated adults of designated genotypes. Scale bars= 2*μ*m. (G) Quantification of nuclei in (A-F). While depleting any single pathway does not have an impact on chromosome morphology in *rad-51;him-5*, impairment of all three pathways increases the occurrence of chromosome fusions. Magenta bars indicate median and interquartile range. Kruskal-Wallis multiple comparisons to *rad-51;him-5* (n.s.: not significant, P>0.06). *rad-51(lg8701);him-5(e1490*) (n=33), *cku-80(ok851);rad-51(lg8701);him-5(e1490*) (n=61), *polq-1(tm2027);rad-51(lg8701);him-5(e1490*) (n=60), *xpf-1(e1487);rad-51(lg8701);him-5(e1490*) (n=37), *polq-1(tm2026),cku-70(tm1524);rad-51(lg8701);him-5(e1490*) (n=61), *xpf-1(e1487); polq-1(tm2026);rad-51(lg8701);him-5(e1490*) (n=50), and *xpf-1(e1487); polq-1(tm2026),cku-70(tm1524);rad-51(lg8701);him-5(e1490*) (n=40).

**Figure S6.**
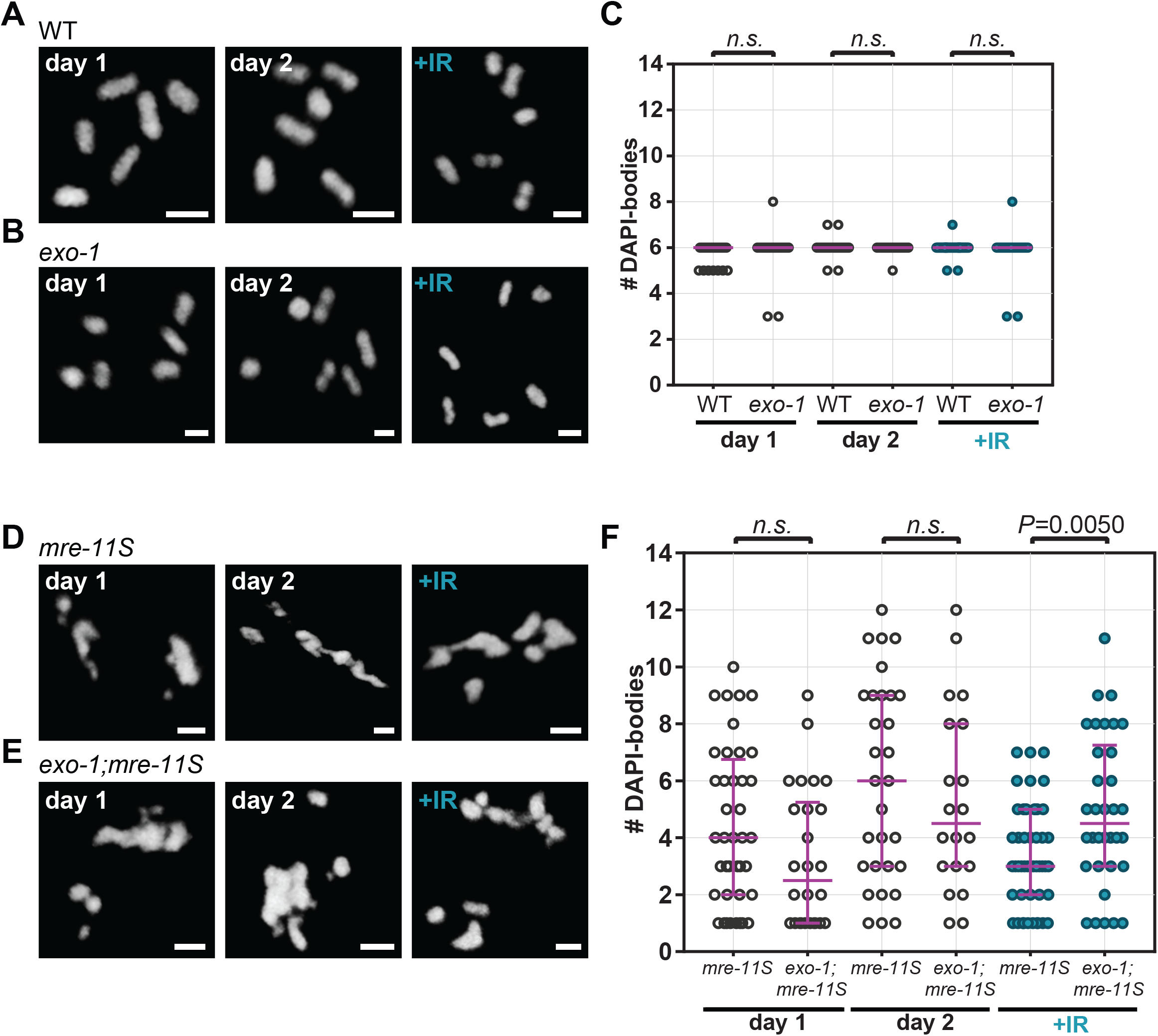
Role of EXO-1 in repair. (A-C) *exo-1* is dispensable for repair of DSBs in an otherwise wild-type background. (A, B) Representative images and (C) Quantification of diakinesis nuclei of one- and two-day-old, adults with and without IR. WT day 1 (n=74), *exo-1(tm1842*) day 1 (n=30), WT day 2 (n=45), *exo-1* day 2 (n=30), WT +IR (n=52), *exo-1* +IR (n=30). Two-tailed Mann-Whitney tests (n.s.: not significant, P>0.48). (D-F) EXO-1 is important to repair IR-induced MRE-11-processed breaks. Upon IR, chromosomes in *mre-11S* mutants have a lower tendency to form fusions if *exo-1* is depleted. (D, E) Representative images and (F) Quantification of DAPI bodies in diakinesis oocytes: *mre-11(iow1*) day 1 (n=40), *exo-1(tm1842);mre-11S* day 1 (n=26), *mre-11S* day 2 (n=30), *exo-1;mre-11S* day 2 (n=20), *mre-11S* +IR (n=47), *exo-1;mre-11S* +IR (n=34). P-values calculated from two-tailed Mann-Whitney tests (n.s.: not significant, P>0.08). Scale bars= 2*μ*m. Dot-plot: Magenta bars indicate median and interquartile range.

**Table S1:** Strains used in this study

| **Genotype** | **Laboratory stock #** |
| --- | --- |
| *brc-2(tm1086) III/hT2 [bli-4(e937) let-?(q782) qIs48] (I;III)* | QP0359 |
| *brc-2(tm1086 III/hT2 [bli-4(e937) let-?(q782) qIs48] (I;III); him-5(e1490) V* | QP1862 |
| *cku-70(tm1524) III* | QP1381 |
| *cku-70(tm1524) III;rad-51(lg8701) IV/ nT1 [unc-?(n754) let-? qIs50] (IV;V).* | QP1671 |
| *cku-80(ok861) III* | QP0842 |
| *cku-80(ok861) III;rad-51(lg8701)/ nT1 [unc-?(n754) let-? qIs50] (IV;V)* | QP1107 |
| *cku-80(ok861);rad-51(lg8701) IV/nT1[unc-?(n754) let-? qIs50] (IV;V);him-5(e1490) V/nT1 unc-?(n754) let-? qIs50] (IV;V)* | QP1073 |
| *dsb-2(me96) II* | QP0938 |
| *dsb-2(me96) II;rad-51(lg8701),rec-8(ok978) IV/nT1[unc-?(n754) let-? qIs50] (IV;V)* | QP1350 |
| *dsb-2(me96) II;rad-51(lg8701) IV/nT1[unc-?(n754) let-? qIs50] (IV;V)* | QP1345 |
| *exo-1(tm1842) III* | QP0901 |
| *exo-1(tm1842) III;mre-11(iow1),him-5(e1490) V/nT1[unc-?(n754) let-? qIs50] (IV;V)* | QP1346 |
| *exo-1(tm1842) III;mre-11(iow1) V/nT1[unc-?(n754) let-? qIs50] (IV;V)* | QP1698 |
| *exo-1(tm1842) III;rad-51(lg8701) IV/nT1[unc-?(n754) let-? qIs50] (IV;V)* | QP1656 |
| *exo-1(tm1842) III;rad-51(lg8701) IV/ nT1[unc-?(n754) let-? qIs50] (IV;V);him-5(e1490) V/ nT1 [unc-?(n754) let-? qIs50] (IV;V)* | QP1657 |
| *exo-1(tm1842) III;rad-51(lg8701) IV/nT1[unc-?(n754) let-? qIs50] (IV;V);mre-11(iow1),him-5(e1490) V/nT1[unc-?(n754) let-? qIs50] (IV;V)* | QP1658 |
| *him-5(e1490) V* | QP0421 |
| *him-5(ea42) V* | QP1398 |
| *him-5(ok1896) V* | QP0432 |
| *mre-11(iow1),him-5(e1490) V/nT1[unc-?(n754) let-? qIs50] (IV;V)* | QP1116 |
| *mre-11(iow1)/nT1[qIs51] (IV;V)* | QP0900 |
| *polq-1(tm2026) III* | QP1244 |
| *polq-1(tm2026),cku-70(tm1524) III;rad-51(lg8701) IV/nT1[unc-?(n754) let-? qIs50] (IV;V)* | QP1592 |
| *polq-1(tm2026),cku-70(tm1524) III;rad-51(lg8701) IV/nT1 [unc-?(n754) let-? qIs50] (IV;V);him-5(e1490) V/ nT1[unc-?(n754) let-? qIs50] (IV;V)* | QP1593 |
| *polq-1(tm2026) III;rad-51(lg8701) IV/nT1[unc-?(n754) let-? qIs50] (IV;V)* | QP1341 |
| *polq-1(tm2026) III;rad-51(lg8701) IV/nT1[unc-?(n754) let-? qIs50] (IV;V);him-5(e1490) V/nT1[unc-?(n754) let-? qIs50] (IV;V)* | QP1547 |
| rad-51(lg8701),rec-8(ok978) IV/nT1[unc-?(n754) let-? qIs50] (IV;V) | QP1347 |
| rad-51(lg8701),rec-8(ok978) IV/nT1[unc-?(n754) let-? qIs50] (IV;V);him-5(e1490) V/nT1[unc-?(n754) let-? qIs50] (IV;V) | QP1351 |
| rad-51(lg8701 IV)/nT1g;mre-11(iow1) V/nT1g | QP1697 |
| rad-51(lg8701) IV/nT1[unc-?(n754) let-? qIs50] | QP1165 |
| rad-51(lg8701) IV/nT1[unc-?(n754) let-? qIs50]; him-5(e1490) V/nT1[unc-?(n754) let-? qIs50] (IV;V) | QP1111 |
| rad-51(lg8701) IV/nT1[unc-?(n754) let-? qIs50]; him-5(ea42) V/nT1[unc-?(n754) let-? qIs50] | QP1411 |
| rad-51(lg8701) IV/nT1[unc-?(n754) let-? qIs50]; him-5(ok1896) V/nT1[unc-?(n754) let-? qIs50] | QP1170 |
| rad-51(lg8701)/nT1[unc-?(n754) let-? qIs50];mre-11(iow1),him-5(e1490)/nT1[unc-?(n754) let-? qIs50] | QP1112 |
| rad-51(lg8701) IV/nT1[unc-?(n754) let-? qIs50];mre-11(iow1) IV/nT1[unc-?(n754) let-? qIs50] | QP1696 |
| rec-8(ok978) IV/nT1[unc-?(n754) let-? qIs50] | QP1297 |
| spo-11(me44),rec-8(ok978) IV/nT1 [qIs51] (IV;V) | QP1348 |
| spo-11(me44) IV/nT1 [qIs51] (IV;V) | QP0963 |
| xpf-1(e1487) II | QP0965 |
| xpf-1(e1487) II;cku-80(ok861) III;rad-51(lg8701 IV)/nT1[unc-?(n754) let-? qIs50];him-5(e1490) V/nT1[unc-?(n754) let-? qIs50] | QP1377 |
| xpf-1(e1487) II;polq-1(tm2026),cku-70(tm1524) III; him-5(e1490) V/nT1[unc-?(n754) let-? qIs50] | QP1595 |
| xpf-1(e1487) II;polq-1(tm2026),cku-70(tm1524) III; rad-51(lg8701) IV/nT1[unc-?(n754) let-? qIs50] | QP1594/QP1700/<br>QP1701 |
| xpf-1(e1487) II;polq-1(tm2026),cku-70(tm1524) III; rad-51(lg8701 IV)/nT1[unc-?(n754) let-? qIs50]; him-5(e1490) V/nT1[unc-?(n754) let-? qIs50] | QP1596/QP1612 |
| xpf-1(e1487) II;polq-1(tm2026) III;rad-51(lg8701) IV/nT1[unc-?(n754) let-? qIs50];him-5(e1490)/nT1[unc-?(n754) let-? qIs50] | QP1376 |
| xpf-1(e1487) II;rad-51(lg8701) IV/nT1[unc-?(n754) let-? qIs50] | QP1167 |
| xpf-1(e1487) II;rad-51(lg8701) IV/nT1[unc-?(n754) let-? qIs50]; him-5(e1490) V/nT1[unc-?(n754) let-? qIs50] | QP1113 |
| rfs-1(ok1372),helq-1(tm2134) III | QP1100 |
| rfs-1(ok1372),helq-1(tm2134) III;him-5(ok1896) V | QP1060 |
| rad-54(ok615) I | QP649 |
| rad-54(ok615) I;him-5(ok1896) V | QP1004 |

**Table S2:**
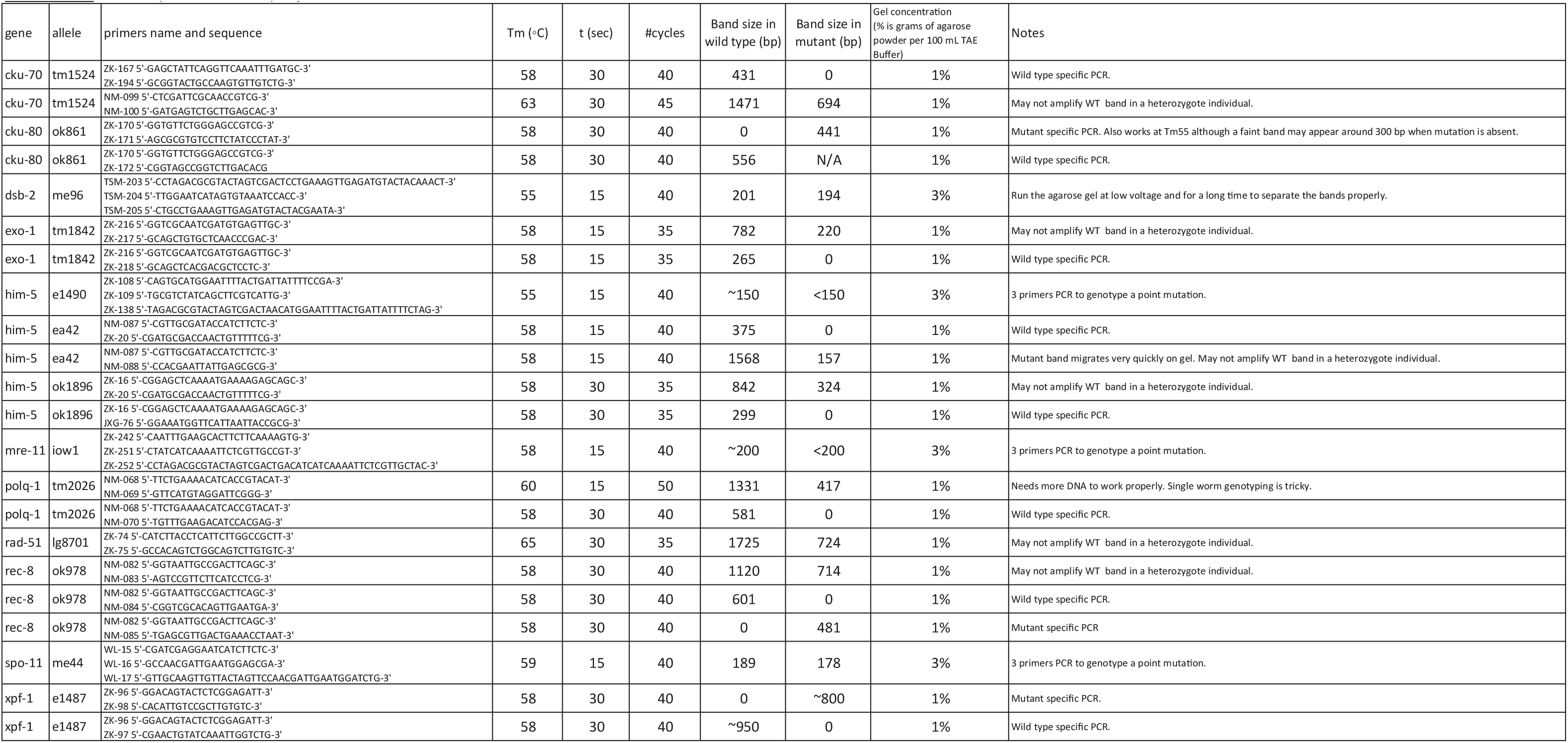

**Table S3:**
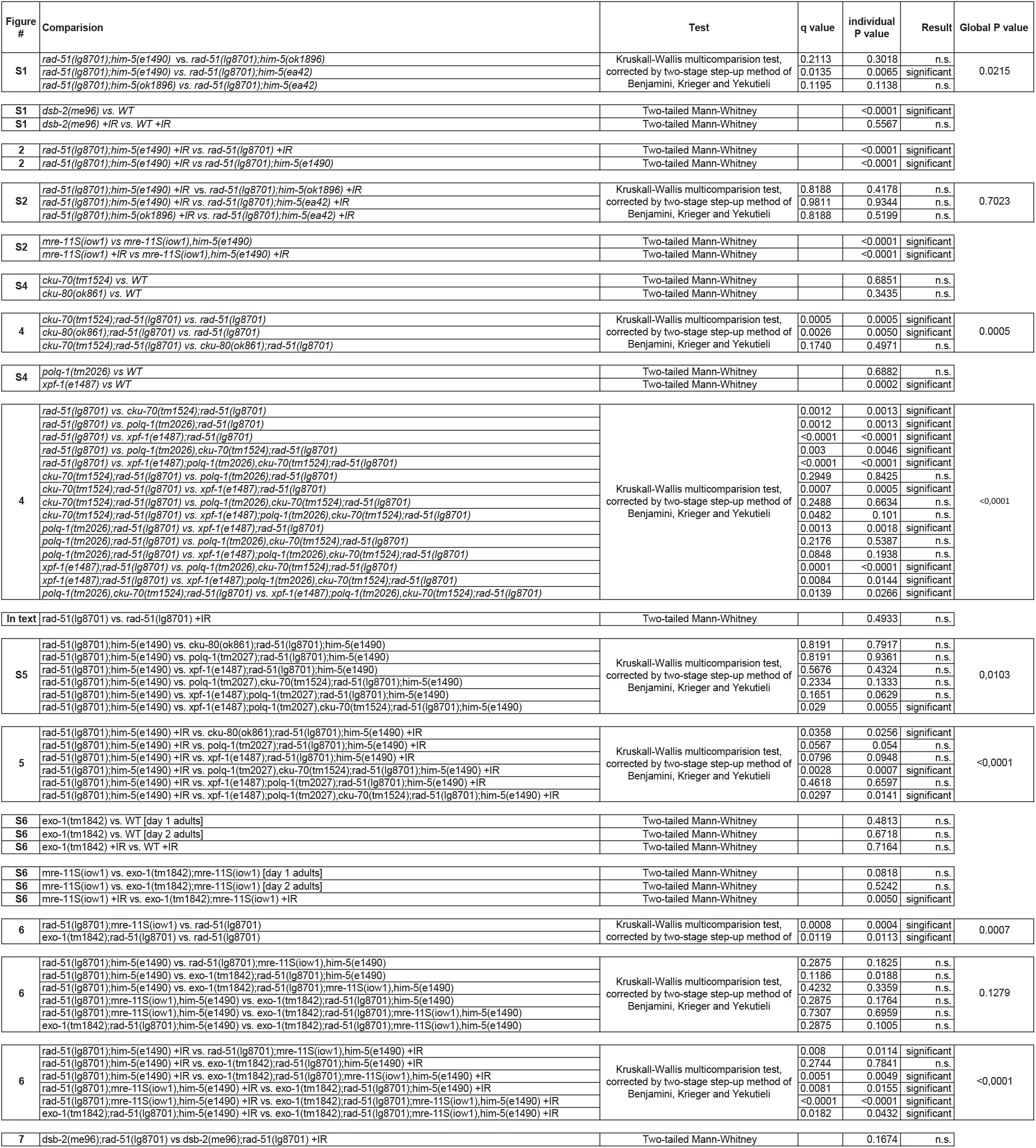

